# Effective and Efficient Neural Networks for Spike Inference from *In Vivo* Calcium Imaging

**DOI:** 10.1101/2021.08.30.458217

**Authors:** Zhanhong Zhou, Hei Matthew Yip, Katya Tsimring, Mriganka Sur, Jacque Pak Kan Ip, Chung Tin

## Abstract

Calcium imaging technique provides the advantages in monitoring large population of neuronal activities simultaneously. However, it lacks the signal quality provided by neural spike recording in traditional electrophysiology. To address this issue, we developed a supervised data-driven approach to extract spike information from calcium signals. We propose the ENS^2^ (<u>e</u>ffective and efficient <u>n</u>eural <u>n</u>etworks for spike inference from calcium signals) system for spike-rate and spike-event predictions using raw calcium inputs based on U-Net deep neural network. When testing on a large, ground truth public database, it consistently outperformed state-of-the-arts algorithms in both spike-rate and spike-event predictions with reduced computational load. We further demonstrated that ENS^2^ can be applied to analyses of orientation selectivity in primary visual cortex neurons. We concluded that it would be a versatile inference system that benefits diverse neuroscience studies.

---

One key to understanding the complex functions of the brain is to simultaneously measure the activity of neurons across different layers and brain areas. Electrophysiological recordings, such as patch-clamp (Hamill et al., 1981; Neher and Sakmann, 1976) and multielectrode extracellular recording (Spira and Hai, 2013), have long been the major method to record neuronal spiking events. These recordings are typically of high temporal resolution and with high signal-to-noise ratio (SNR). However, it is technically challenging with these methods to acquire recordings from a large number of neurons stably *in vivo* (Buzsáki, 2004).

In recent decades, the optical-based two-photon calcium imaging technique has increasingly been used for *in vivo* neuroscience research (de Vries et al., 2020; El-Boustani et al., 2018; Giovannucci et al., 2017; Kerr and Denk, 2008; Knogler et al., 2017; Rikhye and Sur, 2015; Wagner et al., 2017; Wilson et al., 2012). This imaging technique enables simultaneous monitoring of activities of thousands of neurons over a considerable period of time. Moreover, as more effective fluorescent calcium indicators (Akerboom et al., 2012; Bethge et al., 2017; Chen et al., 2013; Dana et al., 2016; Tada et al., 2014) and imaging devices (Denk et al., 1990; Sofroniew et al., 2016; Stosiek et al., 2003) have become available, it is now possible to localize and extract the individual activities of a large number of neurons in various subcellular structures (Grienberger and Konnerth, 2012).

Nevertheless, calcium imaging is only an indirect measurement of neuronal activities. In brief, the concentration of intracellular calcium evoked by neuronal firings undergoes non-linear changes. These fluctuations in calcium are again non-linearly reflected by calcium indicators, whose fluorescent intensities could be imaged. Afterward, the locations of individual neurons or compartments (region of interest, ROI) are identified on images, and the time-varying fluctuations of fluorescence signals in the ROIs are extracted as a surrogate of neuronal activities. Another limitation of calcium imaging is that the signals can commonly have a low SNR (Grienberger and Konnerth, 2012), especially for those recorded in deep brain regions *in vivo* or at low signal conditions. Furthermore, the indicators’ slow temporal dynamics up to hundreds of milliseconds (Chen *et al*., 2013; Kerr et al., 2005) would result in low-pass filtered activities. These indicators come in different types, typically synthetic dyes or genetically encoded calcium indicators (GECIs), and their different dynamics further complicate the task to convert the calcium signal into neuronal signals.

Previous work has shown that spike inference plays a crucial role in interpreting calcium data and dissecting neural circuits (de Vries *et al*., 2020; Kerr and Denk, 2008; Rikhye and Sur, 2015; Wilson *et al*., 2012). In the past decades, researchers have developed various algorithms to recover multiunit neuronal spikes. These algorithms can be generally divided into two major categories: model-based systems (Deneux et al., 2016; Friedrich and Paninski, 2016; Friedrich et al., 2017; Greenberg et al., 2008; Grewe et al., 2010; Jewell et al., 2020; Onativia et al., 2013; Pachitariu et al., 2018; Pnevmatikakis et al., 2013; Pnevmatikakis et al., 2016; Vogelstein et al., 2010; Yaksi and Friedrich, 2006) and data-driven systems (Berens et al., 2018; Hoang et al., 2020; Rupprecht et al., 2021; Sebastian et al., 2021; Theis et al., 2016). In model-based systems, physiologically constrained models were typically built, considering that the calcium signal concentrates with neuronal firings and decays exponentially afterward. With these models, calcium traces could be simulated through estimated spike trains and additive noises. They include systems based on template matching (Greenberg *et al*., 2008; Grewe *et al*., 2010; Onativia *et al*., 2013) (e.g. peeling (Grewe *et al*., 2010)), deconvolution (Friedrich and Paninski, 2016; Friedrich *et al*., 2017; Jewell *et al*., 2020; Pachitariu *et al*., 2018; Vogelstein *et al*., 2010; Yaksi and Friedrich, 2006) (e.g. OASIS (Friedrich and Paninski, 2016)) and Bayes’ theorem (Deneux *et al*., 2016; Pnevmatikakis *et al*., 2013; Pnevmatikakis *et al*., 2016; Vogelstein *et al*., 2010) (e.g. MLspike (Deneux *et al*., 2016)). For example, as one of the state-of-the-art algorithms, MLspike was proposed using a physiologically constrained model and optimized by Maximum a Posteriori (MAP) estimate to infer the most likely spike trains from noisy calcium signals. However, these model-based methods typically require tuning of model parameters for each new recording, either manually or to be estimated by auxiliary algorithms. Moreover, when likelihood optimization is involved, they may become rather computational expensive to use. On the other hand, data-driven systems based on supervised learning also emerged with promising performance. A supervised deep learning algorithm called CASCADE has been reported recently, which delivered high spike inference performance when training data with matched noise levels as the testing neuron were selected for training (Rupprecht *et al*., 2021). Previous data-driven models have faced challenges in validating their generalization ability due to the limited high quality paired data for training (Berens *et al*., 2018; Sebastian *et al*., 2021; Theis *et al*., 2016). An extensive public database of paired data (simultaneously recorded calcium fluorescence signals and electrophysiology ground truths) has been compiled alongside with the development of CASCADE. It has facilitated such data-driven approach for better generalization of the models, although some re-training is still necessary for noise matching (Rupprecht *et al*., 2021). Some other works also use feature extraction and thresholding (Sebastian et al., 2019; Tsutsumi et al., 2015) (e.g. GDspike (Sebastian *et al*., 2019)) to tackle the problem of spike inference.

For inferring unpaired calcium signals from *in vivo* imaging, a calibration-free inference system that could generalize on un-seen recordings with high performance is desirable. In fact, neural networks have shown satisfactory performance in processing bio-signals with severe inter-record variability, including electrocardiogram (Zhai and Tin, 2018; Zhai et al., 2020; Zhou et al., 2021) and electromyography (Zhai et al., 2017). Provided with sufficient amount of paired data, the generalization ability of neural networks makes it a promising approach for inferring spikes from calcium signals. In this work, we performed thorough research on the impact of each component in the neural networks based system on the spike inference tasks. The preferred configurations of network architectures and cost functions were investigated. We conducted additional simulations to address factors in the calcium data that could benefit the performance of deep learning based models for spike inference. These analyses provided useful insights on how to prepare (e.g. record, process, and select) calcium data that will favor future algorithm development, which could help us understand the complex process in the brain. Here, with these research and insights, we developed the ENS^2^ (<u>e</u>ffective and <u>e</u>fficient <u>n</u>eural <u>n</u>etworks for <u>s</u>pike inference from calcium signals) system (Figure 1) with state-of-the-art performance and generalization ability but with lower computational complexity. To further demonstrate the validity of the ENS^2^ system, we deployed the ENS^2^ system on a set of calcium imaging data from primary visual cortex and showed how the spike inference can be applied to the analyses of the experimental data.

**Figure 1.**
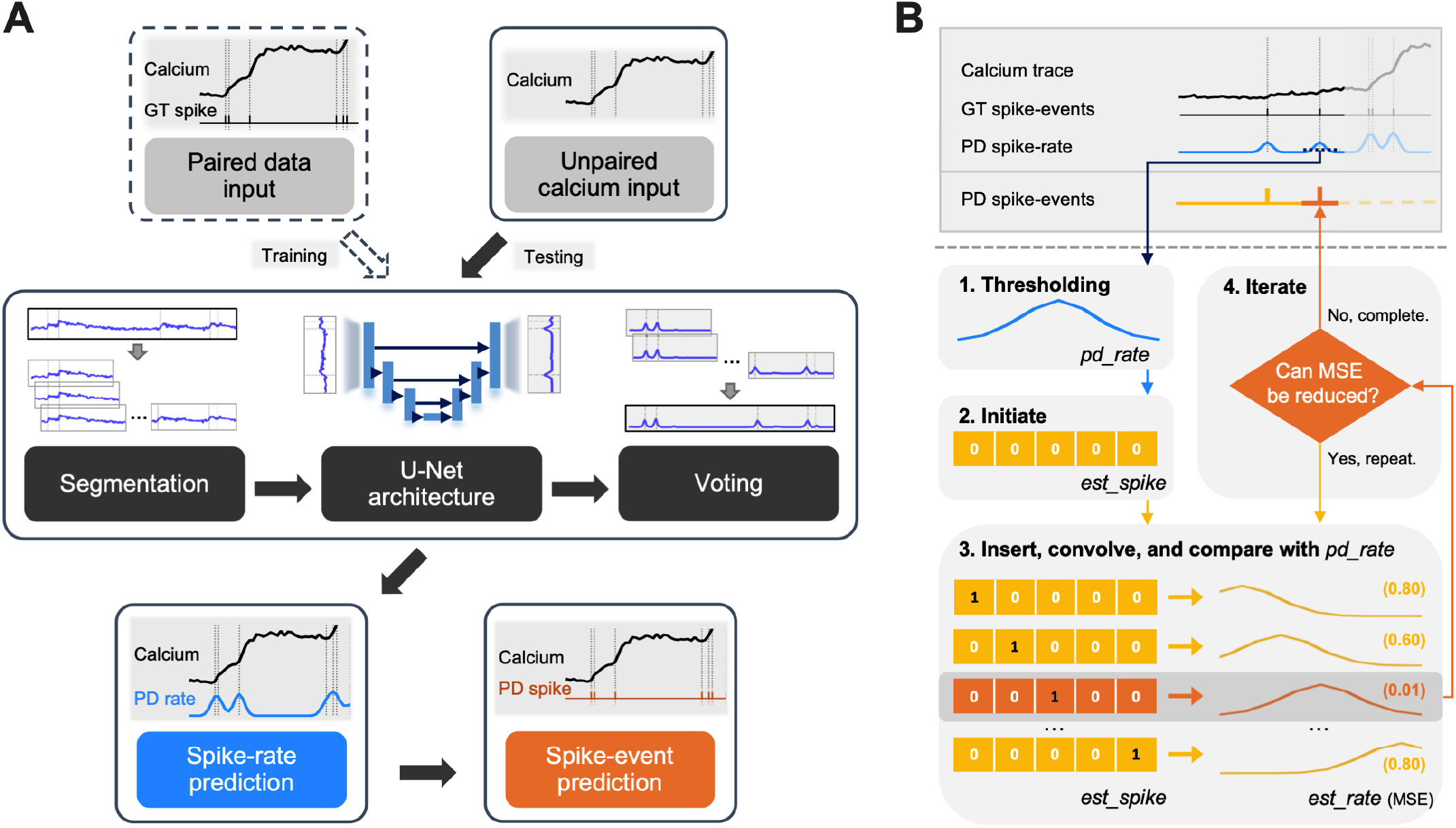
Schematic workflow of the proposed ENS^2^ system. The ENS^2^ system contains a neural network to infer spike-rate from calcium inputs (**A**) and an unsupervised greedy algorithm for estimating spike-events from spike-rate predictions (**B**). The neural network is trained with calcium trace inputs paired with ground truth (GT) spike-events, while it could test on calcium traces alone after training for obtaining predicted (PD) spike-rates. For a given calcium recording, our ENS^2^ system will first predict the corresponding spike-rate with the procedures in (**A**). Inputs are segmented and fed to the U-Net based model, whose spike-rate predictions are gathered through an averaging strategy. Afterward, spike-events are estimated by the four-step algorithm in (**B**). In brief, valid fragments of spike-rate prediction are extracted by thresholding. Then, spike-event is inserted tentatively to approximate the extracted spike-rate fragment. The resultant spike-events sequence that achieves the minimal MSE is regarded as the final spike-events prediction. Details are explained in the Methods section.

## Methods

### Benchmark database

In this study, we used the publicly available datasets containing both calcium imaging signals and simultaneously recorded electrophysiological signals from excitatory neurons (Akerboom *et al*., 2012; Bethge *et al*., 2017; Chen *et al*., 2013; Dana *et al*., 2016; Huang et al., 2020; Khan et al., 2018; Rupprecht *et al*., 2021; Schoenfeld et al., 2021; Tada *et al*., 2014; Theis *et al*., 2016) and inhibitory neurons (Khan *et al*., 2018; Kwan and Dan, 2012; Rupprecht *et al*., 2021). For benchmarking and algorithm development purposes, they were recently compiled by (Rupprecht *et al*., 2021) into an extensive database with 27 datasets. Specifically, we adopted dataset #2 to #27 following (Rupprecht *et al*., 2021) for a fair comparison, and they are labeled as dataset 1 to 26 in this study as shown in Table 1. Specifically, dataset 1 to 20 cover imaging of excitatory neurons from eight different kinds of calcium indicators, a wide range of frame rates (7.7Hz to 500Hz), and various peak firing rates (1.18Hz to 12.67Hz, averaged on each dataset). Over 20 hours of paired ground truth data (calcium signals and spike-events) were recorded from a total of 229 neurons of either mouse or zebrafish brains. On the other hand, dataset 21 to 26 contain inhibitory neurons with much higher peak firing rates (up to 57.31Hz). There is a total of over 15 hours of paired data from 16 *in vivo* and 41 *in vitro* inhibitory neurons.

**Table 1.**
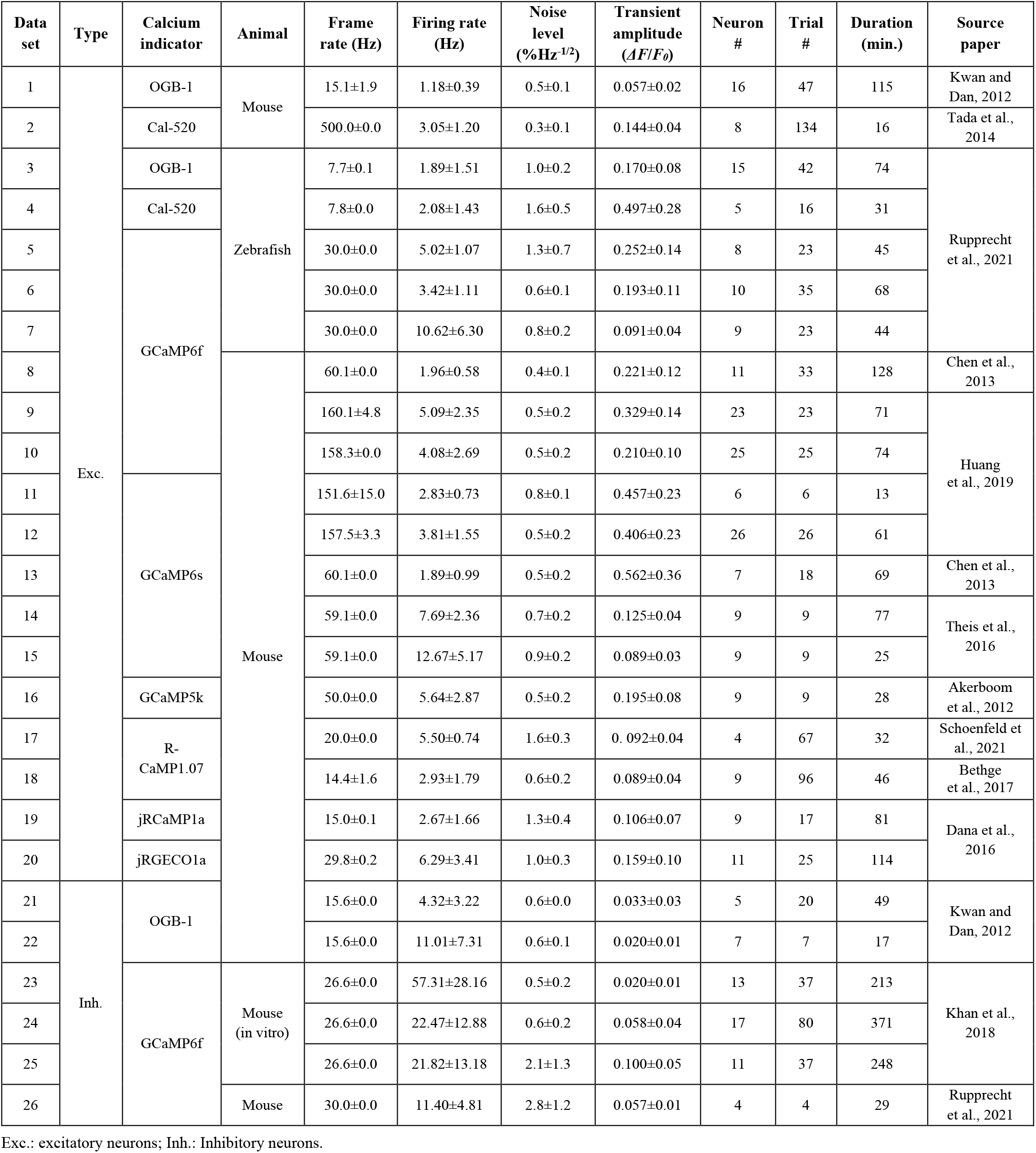
Summary of datasets used in this study (adapted from Rupprecht et al., 2021).

In each dataset, calcium signals are provided as the percentage changes of fluorescence amplitude against baseline (*ΔF*/*F*_*0*_), while individual timestamps label spike-events. We also computed the noise-levels as defined in (Rupprecht *et al*., 2021) and listed them in Table 1. Furthermore, we presented the increase in *ΔF*/*F*_*0*_ induced by one action potential (calcium transient amplitude) for each dataset. The transient amplitude is computed using the averaged calcium kernel, which was extracted from paired ground truth data using the deconvolution function with regularized filter (“deconvreg”) in MATLAB. We computed the approximate instantaneous firing rates of a neuron using a 5 sec. sliding window with steps of 1/60 sec (or 1 data point). The 95% quantile values of these computed firing rates were defined as the peak firing rate of this neuron. The values are shown as mean ± 1 standard deviation in Table 1.

### Data preparation

#### 1) Re-sampling data

To develop and validate the spike inference algorithms, we first re-sampled the input data (both training set and testing set) of different frame rates to the same sampling rates. In this work, we referred to the original frequencies where calcium signals were captured as *frame rates*, and the re-sampled frequencies as *sampling rates*. Given that most of the datasets were captured with frame rates not higher than 60Hz (Table 1), we re-sampled all calcium signals to 60Hz. All the inference systems were then benchmarked under this same sampling rate. We also tested our system under 7.5Hz as suggested by CASCADE (Rupprecht *et al*., 2021). The re-sampling is performed with the “resample” function of SciPy. The impact of sampling rates on inference results is discussed in this work.

#### 2) Pre-processing of inputs

We used the original *ΔF*/*F*_*0*_ calcium inputs for our system (where only re-sampling is performed). The training target (expected outputs from the systems) are prepared as below. For a pre-defined sampling rate (e.g. 60Hz), raw timestamps of ground truth spike (spike-events) are re-allocated into their corresponding time bins. We can then compute the sequence of spike counts by counting the total firing events in each time bin. The sequences are then smoothed with Gaussian filters to facilitate gradient descent. The smoothing window size τ for the Gaussian kernels was set to 25ms, which produces the optimal spike-event predictions with high temporal resolution in general. The selection of smoothing window size for deep learning based systems is also carefully studied. The convolved spike counts are denoted as “spike-rate” in this work.

#### 3) Data segmentation

To train the neural networks properly, paired sequences of calcium signals and spike-rates were segmented with a moving step of 1 data point (Figure 1A). The length of each segment was set to 96 data points. In the case of sampling rate of 60Hz, a total of ∼4 million segments of paired data were obtained for training.

### Network architectures

We tested three different architectures of neural networks to evaluate their effectiveness in the spike inference task. They are U-Net (Ronneberger et al., 2015), Le-Net (Lecun et al., 1998), and FC-Net (fully-connected network), respectively. These existing models were modified for 1-D calcium signal inputs. The network architectures are summarized in Table 2.

**Table 2.**
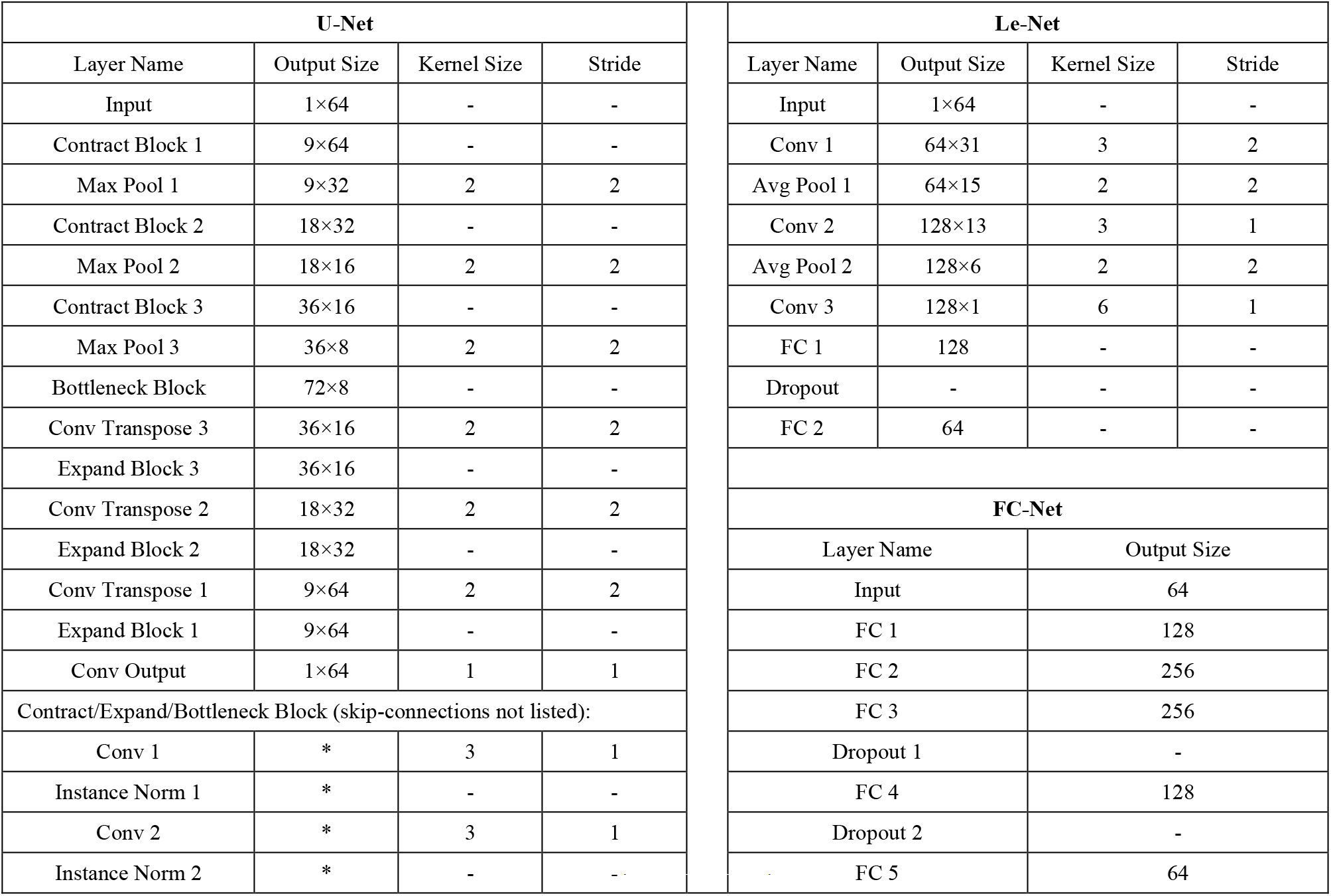
Summary of network architectures.

The U-Net used in this study contains three contracting blocks and expanding blocks. On one hand, the input information from contracting blocks passes through the bottleneck block to the expanding blocks. On the other hand, skip-connections from contracting blocks to the corresponding expanding blocks allow direct and localized information flows (Ronneberger *et al*., 2015). Within each contracting/expanding block and the bottleneck block, two convolution layers with 3-sized kernels are deployed. Instead of batch normalization, we used instance normalization (Ulyanov et al., 2016) for regularization, since calcium signals with various dynamics may co-exist in a same batch of data. We observed that this regularization helped in model convergence. The Le-Net consists of three convolution layers with kernel sizes of 3, 3, and 6, respectively. Average pooling layers with 2-sized kernels are applied between the three convolution layers. A dropout layer (Srivastava et al., 2014) is included before the output layer for regularization. A typical fully-connected network with four hidden layers and two dropout layers is adopted as FC-Net. All three networks are designed to take 96-sized calcium signal inputs, and output 96-sized spike-rate vectors in a sequence-to-sequence translation manner. During prediction, given that the input data are segmented with steps of 1 data point, each time point is indeed predicted for up to 96 times separately from its adjacent segments. We, thus, were able to average these predictions for a robust final spike-rate output of each time point (Figure 1A). Spike-event output could then be estimated from this final spike-rate sequence as introduced below.

We kept all three networks to have similar numbers (under 150k) of trainable parameters for comparison (Figure 5A). They are all randomly initiated to have zero means and standard deviations of 0.02. Leaky ReLU (rectified linear units) with slopes of 0.2 is used as activation functions for all layers except for the output layers, where ReLU is used for non-negative spike-rate prediction.

### Loss functions and optimization

For each type of network, we optimized them with three different loss functions, respectively, for comparison. First, mean square error (MSE) loss is used, which is one of the most commonly used loss functions applicable to a wide variety of machine learning tasks. The models are expected to minimize the MSE between predicted spike-rates and ground truth spike-rates, penalizing the prediction both in time and amplitudes. The loss function is as follows:

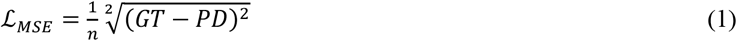

where *n* is the number of segments in a batch. GT and PD stand for ground truth and prediction, respectively. In addition, we also used Pearson correlation coefficient (Corr) and van Rossum distance (vRD) (van Rossum, 2001), as loss functions:

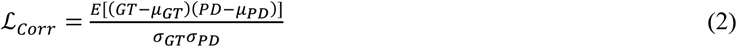

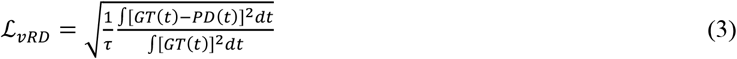

where *μ* and *σ* are the mean and standard deviation, respectively, and τ is a constant normalizing factor (see the evaluation section below). Note that Corr and vRD are also used as the metrics for evaluating the performance of the models (See below). As such, we would expect that models that use ℒ_*Corr*_ (or ℒ_*νRD*_) should have the optimal performance when measured with Corr (or vRD). A major difference between ℒ_*MSE*_ and ℒ_*νRD*_ is that the ℒ_*νRD*_ normalizes each batch of samples by the total numbers of GT firing events. As such, the optimization through ℒ_*νRD*_ is less dependent on the firing rates of the training data, but is slightly more computationally expensive to use than ℒ_*MSE*_. We will compare their resultant inference performance in further detail.

The Adam optimizer (Kingma and Ba, 2015) with a default learning rate of 1e-3 is used for all models. Each model is allowed to update for a maximum of 5000 iterations. In each iteration, a batch of 1024 paired segments is drawn randomly and fed to the model for training. The training losses are noted, and early-stopping is introduced when the losses do not improve in the past 500 interactions (*patience* = 500). Under these criteria, we observed that most models completed the trainings within 3000 iterations. The resultant models are then ready for prediction. Other details of hyper-parameters and operational environment are summarized in Table 3.

**Table 3.** Summary of hyper-parameters and operational environment.

|  |  |
| --- | --- |
| Language | Python 3.6.12 |
| Scientific computing | Numpy 1.19.2, Scipy 1.5.2 |
| Deep learning toolbox | PyTorch 1.7.1 |
| Data processing | MATLAB 2020b |
| CUDA version | 10.1 |
| CUDNN version | 7.6 |
| Dropout rate | 50% |
| Weight initialization | $\mathcal{N}(0,0.02)$ |
| Optimizer | Adam |
| Learning rate | 1e-3 |
| Batch size | 1024 |

### Estimation of spike-events from spike-rate predictions

To reliably convert the spike-rates output by the neural networks to spike-event predictions, we propose an unsupervised greedy algorithm that is simple and straight-forward (Figure 1B). The workflow is briefly introduced here.

Step 1: Fragments of spike-rate predictions (*pd_rates*) with non-zero spike-rate are identified by thresholding the spike-rate sequence output with an epsilon value. We do not use zero threshold to avoid including any fragment with overly low peak amplitude (i.e. those showing extremely small spiking probabilities or background noise), where no spike should be estimated.

Step 2: For each *pd_rate* of length *L* (in terms of number of data points), we initialize a zero-filled vector (*est_spike*) with *L* bins.

Step 3: One spike is assigned to any one bin in *est_spike* at one time and convolve it into a spike-rate vector (*est_rate*) in a same way as we have described above. Then the MSE between the resultant *est_rate* and *pd_rate* is calculated. This step is implemented parallelly for all *L* bins to determine the most suitable bin (i.e. with the smallest MSE) for assigning the spike.

We then repeat step 3 to assign another spike each time to the most suitable bin in a greedy manner, until the MSE would no longer be reduced by adding a spike to any location in *est_spike*. Then the updated *est_spike* is regarded as the final estimation of spike events for the concerned *pd_rate* fragments. The timestamp of a spike is defined as the center time of the corresponding bin within *est_spike*. If multiple spikes are predicted in the same bin, the same timestamp is repeated accordingly.

For a spike-rate sequence output with *N* fragments of *pd_rates*, this algorithm executes in *O*(*N*×*L*×*k*) time, where *k* is the maximum number of spikes in any one bin. In practice, considering the typically slow dynamics of calcium signals and relatively low firing rates of neurons imaged, this estimation method operates in linear time in proportional to the duration of recordings. We have validated our system with this spike-events estimation algorithm against several existing studies, including MLspike (Deneux *et al*., 2016), OASIS (Friedrich *et al*., 2017), and CASCADE (Rupprecht *et al*., 2021) with its Monte-Carlo importance sampling based spike-events estimation algorithm.

### Evaluation

How to reliably assess the performance of the spike inference tasks remains an open topic, where a single evaluation metric could be biased in certain aspects (Berens *et al*., 2018; Rupprecht *et al*., 2021; Sebastian *et al*., 2021; Theis *et al*., 2016). In this regard, recent studies proposed to employ multiple metrics to supplement each other (Berens *et al*., 2018; Deneux *et al*., 2016; Hoang *et al*., 2020; Rupprecht *et al*., 2021; Sebastian *et al*., 2021; Theis *et al*., 2016). In this work, we used four metrics to examine spike-rates prediction and two others for spike-events prediction.

Firstly, Pearson correlation coefficient (Corr) is used as the primary metric for comparing similarities of spike rates as follow:

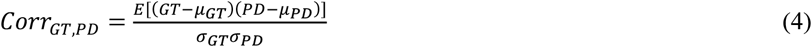

where GT and PD stand for ground truth and prediction, respectively. Secondly, we use the van Rossum distance (vRD) (van Rossum, 2001) for the evaluation of spike rates prediction:

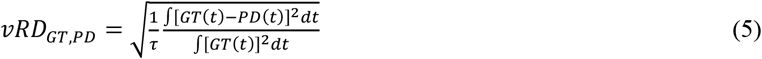

where the time constant *τ* is the normalizing factor (smoothing window size) for smoothing spike-events into spike-rates (e.g., *τ* = 0.025*s* for our proposed system). Moreover, Error and Bias proposed in (Rupprecht *et al*., 2021) are also used to evaluate spike-rates:

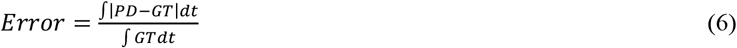

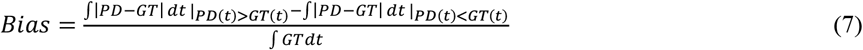

On the other hand, for measuring spike-event prediction, we adopt the Victor-Purpura distance (VPD) (Victor and Purpura, 1996). It is defined as the minimal cost to transform the PD spike-events to the GT spike-events. The cost for either inserting or deleting a spike equals 1, while shifting a spike by Δ*t* costs *q*|Δ*t*|. We use the default value *q* = 1 in this work. To make comparison across different datasets, we present the VPD as the minimal total cost divided by the total number of GT spikes.

Lastly, we compute the error rate (ER) as below (Deneux *et al*., 2016; Éltes et al., 2019; Hoang *et al*., 2020), which measures the F_1_ score of the predicted spike-events:

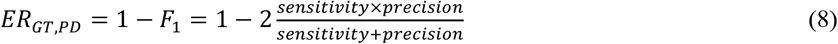

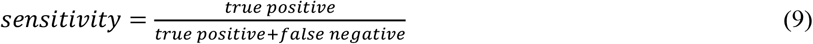

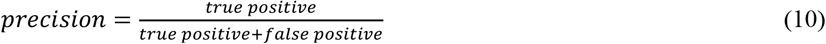

The GT spike-events and PD spike-events are matched based on their VPD. Here, a spike is said to be correctly predicted if it co-exists with its real counterpart within a time window of 50ms (defined as the ER window size). This time window is one order smaller than that used in previous study (Deneux *et al*., 2016), suggesting a much more stringent assessment of model performance in this study. We also examined the effect of ER window sizes in this work.

### Data analysis and statistical testing

During our benchmarks, we recorded the above metrics for each neuron in the testing dataset. To compare among different system configurations and models, we show their medians with 95% confidence intervals (Figure 2A-D, Figure 3A-D, Figure 5B-D, Figure 7B, Figure 10G, Supplementary Figure 1A-D, Supplementary Figure 2A-F, Supplementary Figure 3, Supplementary Figure 5B-G, Supplementary Figure 6) among all neurons from all testing datasets. We performed Shapiro-Wilk test before subsequent statistical analyses, and all did not pass normality tests. We then used Friedman test with Dunn’s multiple comparison for statistical analyses when the number of paired groups are larger than two (Figure 2A-D, Figure 3A-D, Figure 5C, Figure 7B, Supplementary Figure 1A-D, Supplementary Figure 3), and two-sided Wilcoxon signed-rank test otherwise (Figure 5B&D, Figure 9B-E, Supplementary Figure 1J-K, Supplementary Figure 5B-G, Supplementary Figure 6). When testing on the benchmark dataset (Figure 2A-D, Figure 3A-D, Figure 5B-D, Figure 10G, Supplementary Figure 1A-D, Supplementary Figure 2A-F, Supplementary Figure 3, Supplementary Figure 5B-G, Supplementary Figure 6), the performance resulted from two competing models on the same testing neuron is regarded as a paired sample. For the visual stimulating experiment (Figure 7B), the selectivity indexes obtained from *ΔF*/*F*_*0*_, spike-rate, or spike-event on a same neuron are regarded as a paired sample. We reported the significance with p values (^*^p<0.05, ^**^p<0.01, ^***^p<0.001, ^****^p<0.0001, ns: not significant).

**Figure 2.**
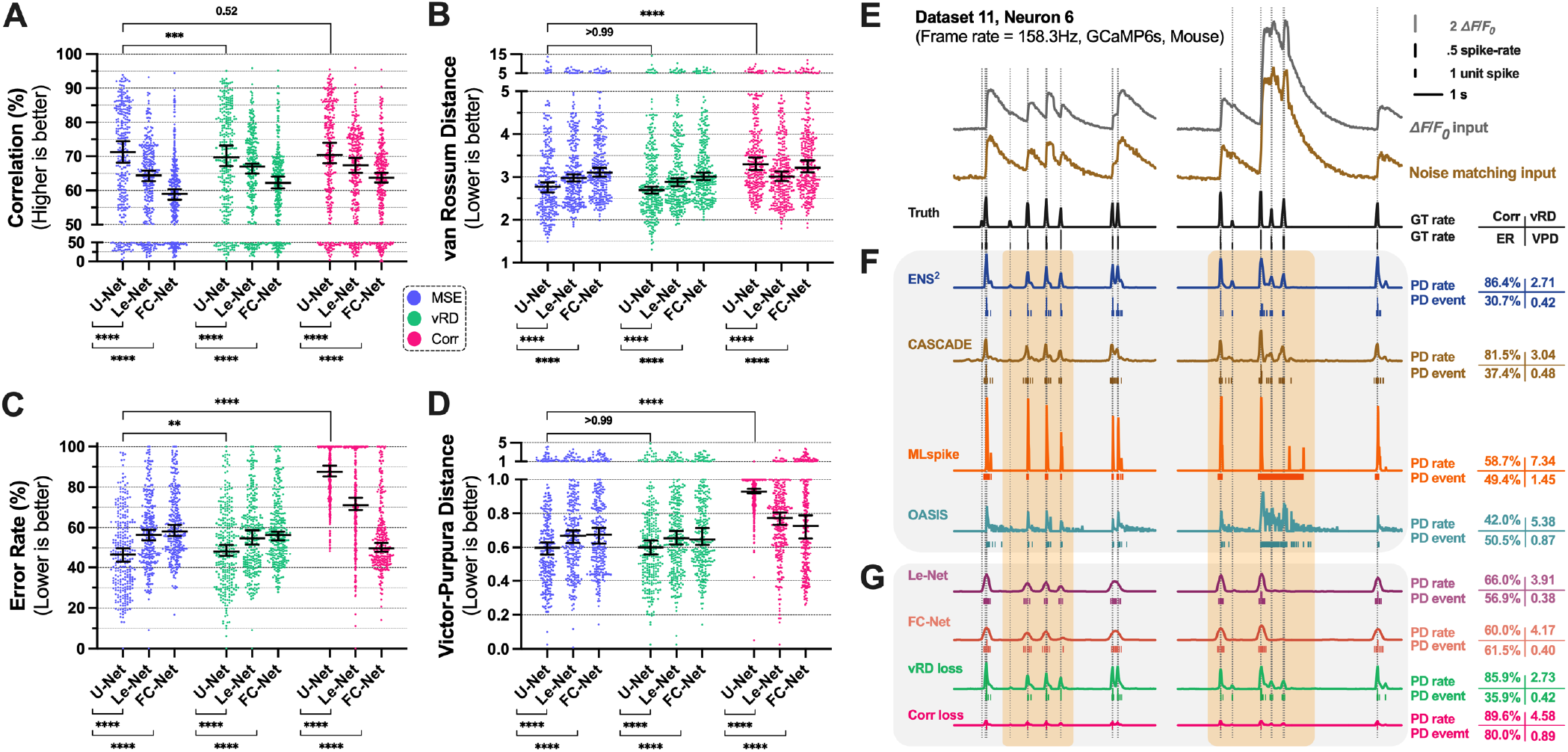
Performance comparison among various model architectures and loss functions. Performances of various system configurations measured in (**A**) correlation, (**B**) van Rossum distance, (**C)** error rate, and **(D**) Victor-Purpura distance, respectively. Generally, the configuration of U-Net with MSE loss provides the best overall performance for both spike-rate prediction and spike-event prediction. (**E**) Examples of different (pre-processed) calcium inputs (colored), paired with ground truth spike-rate and spike-events (black). (**F**) Examples of spike-rates and spike-events predicted by our proposed ENS^2^ method and state-of-the-art methods. (**G**) Examples of spike-rates and spike-events predicted by our proposed method with different configurations. All error bars in this work present medians with 95% confidence intervals. Asterisks show the results of Friedman test with Dunn’s multiple comparisons between the indicated configurations (see Methods, n=286, ^*^p<0.05, ^**^p<0.01, ^***^p<0.001, ^****^p<0.0001). Colored dots in **A**-**D** represent measured performance of each neuron. Metrics shown in **F**-**G** measure the performance on the corresponding neuron. Orange shaded areas represent regions of interest where discrepancies in predictions are observed among different methods. These conventions are consistent for the remaining figures.

**Figure 3.**
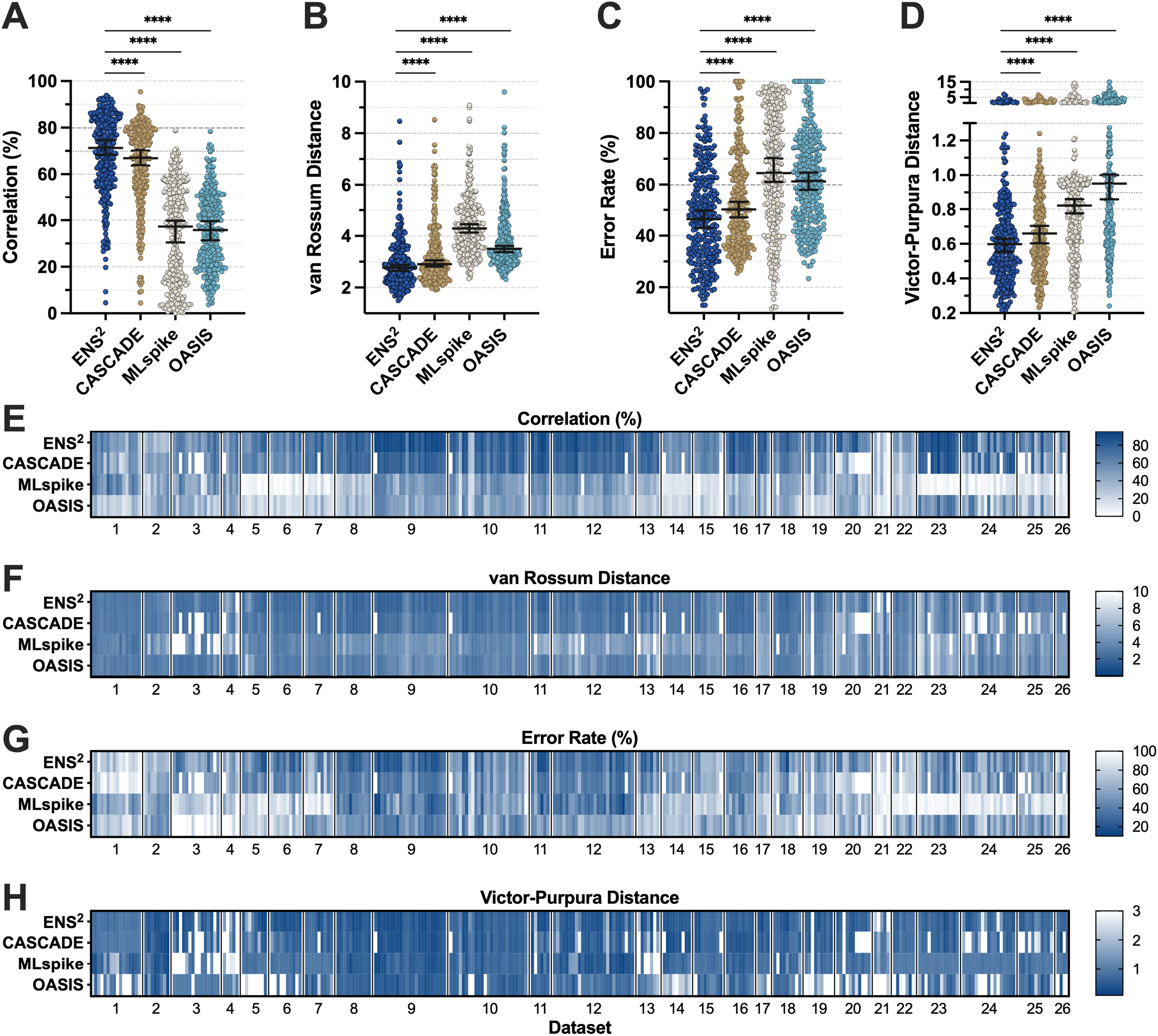
Performance comparison of our proposed ENS^2^ system against state-of-the-art algorithms. Neuron-wise performance is measured in (**A**) correlation, (**B**) van Rossum distance, (**C**) error rate, and (**D**) Victor-Purpura distance, respectively. Inference performance on each neuron in all datasets is summarized in (**E**-**H**) heatmaps. Colored circles in **A**-**D** denote the performance for each individual neuron. Error bars present medians with 95% confidence intervals. Asterisks show the results of Friedman test with Dunn’s multiple comparisons between the indicated systems.

### Implementation of algorithms

As described above, this study involves configurations from three types of neural network models and three loss functions, resulting in a total of 9 configurations of models. The best performing system as evaluated by the six metrics described in the Method section was then benchmarked against the state-of-the-art algorithms, including the data-driven method, CASCADE (Rupprecht *et al*., 2021) and the model-based methods, MLspike (Deneux *et al*., 2016) and OASIS (Friedrich *et al*., 2017).

Our simulations followed the leave-one-dataset-out protocol. For example, when benchmarking on excitatory neurons, each time the model was first trained on 19 datasets and tested on the remaining one. This was repeated 20 times such that all the 20 datasets (#1-#20) with excitatory neurons were tested respectively. The procedure is similar for the 6 datasets (#21-#26) with inhibitory neurons. We then recorded the neuron-wise performance in all 26 datasets.

For the CASCADE algorithm, we follow the training protocol as described (Rupprecht *et al*., 2021). For each dataset, the “noise matching inputs” were obtained from CASCADE to reproduce results with the algorithm (e.g. artificial noise is added to the 19 training datasets to match the noise-level of the testing excitatory neurons, and vice versa for inhibitory neurons). Five identical models were trained separately for 10 epochs. The averaged outputs of these five models were regarded as the final spike-rate predictions. When testing under 60Hz sampling rate, the smoothing kernel size was reduced from 0.2s to 0.025s. Spike-event predictions are estimated using a Monte-Carlo importance sampling based algorithm in CASCADE (Rupprecht *et al*., 2021). All hyperparameters are kept intact as in the CASCADE algorithms.

For the MLspike algorithm (Deneux *et al*., 2016), original *ΔF*/*F*_*0*_ calcium inputs are used. For datasets with OGB synthetic dyes, we set saturation *γ* = 0.091. For datasets with GECIs, we used the full physiological model version of MLspike with parameters modeling saturation, Hill exponent, c0, and rise time as described in Supplementary Figure 6a of (Deneux *et al*., 2016). For the remaining datasets, we used the polynomial nonlinearity modeling in MLspike with coefficient [*p*_2_, *p*_3_] = [1.0, 0.0]. The values of the model parameters, *A* (transient amplitude), *τ* (calcium decay time constant), and σ (noise amplitude) were obtained using its built-in auto-calibration algorithm, since manual calibration on fresh recordings without ground truth is also challenging in actual application (see Discussion). In case that the auto-calibration failed, we supplied the parameters of *A* and *τ* manually (following the methods shown in Supplementary Figure 6a of (Deneux *et al*., 2016)). For a fair comparison, in addition to the spike-event outputs from MLspike, we also obtain its native *spikest_prob* output for evaluating its performance in spike-rate inference.

For the OASIS algorithm (Friedrich *et al*., 2017), we used the L_1_-regularized version by calling the *deconvolve* function. Original *ΔF*/*F*_*0*_ calcium traces were fed as inputs. We set *optimize* _*g* = 5 to auto-calibrate the parameter *g*. The hard thresholds for obtaining discrete spike-event outputs were computed as 55% (slightly over one half as described in OASIS) of the real calcium transient amplitude of each neuron. Note that such amplitudes of the testing recording might be unavailable in actual application without ground truth paired data (see Discussion).

### In vivo experiments

#### 1) Animal preparation and surgery

Ai148 mice (Jackson Lab strain #030328) were crossed with CamKII-cre mice to express GCaMP6s in excitatory neurons. Animal surgery was described previously (El-Boustani *et al*., 2018). In brief, postnatal day (P) 25 mice were anaesthetized with 3% isoflurane and confined by stereotaxic frame. Scalp was sterilized and removed for the cranial window surgery. Skull above the left binocular visual cortex (Figure 6A3) was replaced by a 3mm/5mm stacked circular glass coverslip to ensure transparency for imaging. A tailor-made head-plate was fixed on the skull with Metabond adhesive cement. All procedures performed on mice were conducted under protocols approved by the Massachusetts Institute of Technology’s Animal Care and Use Committee and conformed to US National Institutes of Health (NIH) guidelines.

#### 2) Two-photon calcium imaging

The imaging process was performed by two-photon system with awake and head-fixed mice. The mice were allowed to recover for at least 3 days after the craniotomy, followed by a habituation on head-fixation. Prairie Ultima with a Spectra Physics Mai-Tai Deep See laser two-photon system (Prairie Technologies) was used for imaging. 20x Olympus objective lens was used for functional imaging. The calcium signals of neurons from layer 2/3 were visualized at 920nm laser wavelength with acquisition frame rate around 7.6Hz (averaging from 4 consecutive frames around 30Hz).

#### 3) Visual stimulation protocols

The visual stimulus was delivered from Psychtoolbox-3. A computer was connected to a 10-inch 1080p LCD monitor for display. Drifting gratings were used to stimulate the visual cortex (Figure 6A2-3). Each trial of display lasts for 10 seconds and repeated for 20 trials per stimulus, resulting in imaging session of 200 seconds. At the beginning of each trial, 6 seconds of grey screen is presented as blank, followed by 4 seconds of grating stimuli (ON period). 8 directions were presented from 0 to 315 degrees to the horizontal, while each direction was displayed for 0.5 second and started with another direction of 45 degrees increment (Figure 6A3).

#### 4) Data processing and analysis

The recorded images stacks were processed in Fiji (ImageJ version 1.53c) before data extraction. The slices from contralateral (*contra*) and ipsilateral (*ipsi*) recordings (with respect to the visual stimuli) were combined and stacked by maximum intensity Z-projection to concatenate into a single movie. The motion artifact was minimal with plugin ‘Template Matching’ by recognizing and aligning blood vessels over large region within slices. The Z-projected slice was kept for alignment only. The aligned movies were imported to Suite2P (Pachitariu et al., 2017) (version 0.10.1, https://github.com/MouseLand/suite2p) for neuron segmentation. The following parameters were changed from the default: tau of 0.75, denoise of 1, diameter of 9, anatomical only of 1, maximum iterations of 1, frames per second of 5 and 191 minimum neuropil pixels. The regions of interest (ROIs) and corresponding cells’ activities were detected and saved in.mat file for further processing in MATLAB.

The ROIs of cell were identified with Suite2P-generated file, *iscell*, and the non-cell components were removed. Z-score and the relative change in fluorescent (*ΔF*/*F*_*0*_, where *F*_*0*_ was the fluorescence baseline of that ROI) was calculated. A two-step approach was used to further select the visually responsive neuron, which showed significant activities to specific direction(s). First, the *ΔF*/*F*_*0*_ of each direction was averaged within trial and compared with the corresponding *ΔF*/*F*_*0*_ of 4^th^ – 6^th^ second of blank with two-sided t-test at 5% significance level. Afterwards, the neurons with significant difference at any direction were searched for exactly 3 consecutive frames of ON period with z-score > 3. The neurons passed both criteria are regarded as visually responsive. Only these visually responsive neurons were tested with the spike inference algorithm.

#### 5) Calculation of tuning curves and selectivity indexes

For each trial in the recording of responsive neurons, mean responses within each 0.5sec stimulus window are taken (e.g. mean *ΔF*/*F*_*0*_ for calcium traces and mean firing rate for spike-events predictions), producing 8 different mean response values. We also compute the background responses using the averaged activities within the 6sec resting window. Then, the resultant 8 mean response values are subtracted from their corresponding background responses in each trial. The final tuning curves are obtained by averaging these mean responses across 20 trials.

We adopted the OSI (orientation selectivity index) and DSI (direction selectivity index) as defined in a previous study (Mazurek et al., 2014) to quantify the tuning curves and neuronal selectivity further,

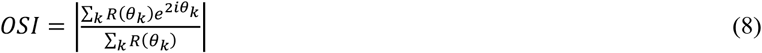

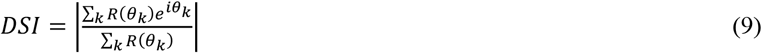

where *R (θ* _*k*_) is the response to stimulus orientation at angle *θ*_*k*_, and *k* = 8 is the total number of stimulus angles. Before computing OSI/DSI, if any value in a tuning curve is below zero, we up-shift the whole tuning curve to keep it non-negative for calculation of OSI and DSI. Supplementary Figure 7 shows several representative examples of how different patterns in tuning curves could affect OSI/DSI quantitatively. The OSI/DSI of the neuron population were quantified with bar plots showing their medians and 95% confidence intervals, and statistically compared (Figure 7B).

## Results

### U-Net and MSE loss achieve the best overall performance in spike inferring tasks

We evaluated all potential configurations of our models across all 26 datasets with the leave-one-dataset-out protocol (see Methods), such that the testing data was not “seen” in the training data. The results are presented for each neuron from the testing datasets (Figure 2A-D). All datasets were resampled to a frame rate of 60Hz for benchmark (see Methods).

First, we compared three architectures of neural networks (U-Net, Le-Net, and FC-Net, see Methods). As shown in Figure 2A-D, U-Net delivered the best overall performance in all four evaluation metrics against either Le-Net or FC-Net (p<0.0001 for all cases, when MSE or vRD loss function was used). Next, we assessed how different loss functions would affect the performance of our models. Here, MSE, vRD and Corr were used as the loss functions, respectively. The vRD and Corr were used as loss function to test if they would favor spike-rate prediction since they are evaluated by vRD and Corr. Nevertheless, our results show that MSE loss appeared to be the better choice in general. When measured in Corr (Figure 2A), using MSE loss showed similar performance with using Corr loss (p=0.52), and obtained higher Corr compared to vRD loss (p=0.0006). When measured in vRD (Figure 2B), using MSE loss also showed similar performance with using vRD loss (p>0.99), and obtained lower vRD compared to when using Corr loss (p<0.0001). For spike-event predictions, using MSE loss again gained advantages over using vRD loss in ER measurement (p=0.0032, Figure 2C), and comparable performance in VPD measurement (p>=0.99, Figure 2D). Note that when Corr loss was used, the performance of all three models significantly compromised, except when measured in Corr. The degradation in performance was most prominent in spike event predictions (measured in ER and VPD (Fig. 2C-D)). We believe that the major reason is that the Corr is a scale-free measurement, and Corr loss function fails to differentiate predictions of different amplitudes (but only different temporal patterns). As shown in Figure 2G, the prediction obtained with models using Corr loss tended to have much lower spike-rate (in amplitude) as compared to other configurations (Figure 2E-G). As a consequence, spike-event could not be reliably estimated from the predicted spike-rate due to low signal-to-noise ratio. Putting the above results together, we took U-Net and MSE loss function in our proposed ENS^2^ as it achieved the best overall performance in the benchmark.

To show that the difference in performance resulted from the model configurations rather than specific hyper-parameter settings, we repeated the simulation with U-Net and MSE loss with various hyper-parameters (Supplementary Figure 2). The filled bars represent the default hyper-parameter combination used in this study as described in the Method section. Regardless, we showed that they all had little effect on the final performance. The Corr approached 70% and vRD remained less than 3 in all cases. The VPD and ER were around 0.6 and 50%, respectively.

We also proved that our models were trained adequately with our early-stopping criteria (see Method). Supplementary Figure 2G illustrates the MSE training losses for all 20 datasets with excitatory neurons (red) and all 6 datasets with inhibitory neurons (blue). The losses decreased with more iterations generally and stabilized sufficiently as the training stops. Moreover, Supplementary Figure 2 demonstrates that the patience of iterations (see Method) before early-stopping had little influence on performance. Together, we proved that our models have been trained and regularized sufficiently by iterating over only thousands of batches of data to avoid over-fitting.

### Comparison to state-of-the-art studies

Based on our investigations above, we selected the configuration of U-Net and MSE loss as our proposed ENS^2^ system, which takes original *ΔF*/*F*_*0*_ signals as inputs. We took this further to compare it with three representative state-of-the-art studies: CASCADE (Rupprecht *et al*., 2021), MLspike (Deneux *et al*., 2016), and OASIS (Friedrich *et al*., 2017). We selected CASCADE and MLspike as they are the among the top performing systems within the two major categories: data-driven systems and model-based systems, respectively. Moreover, both of them have already shown surpassing performance over previous methods using various datasets and evaluation metrics in their studies. On the other hand, OASIS is one of the most representative methods based on deconvolution, and has been implemented in Suite2P (Pachitariu *et al*., 2017) and CalmAn (Giovannucci et al., 2019) for widely experimental usages. Results are summarized in Figure 3.

We first benchmarked their performance across all 26 datasets (see Methods). Figure 3A-D shows that the data-driven systems (i.e. our proposed ENS^2^ and CASCADE) generally performed better than the model-based system (e.g. p<0.0001 for all cases of ENS^2^ vs. MLspike/OASIS), for both spike-rate (Corr and vRD) and spike-event (VPD and ER) predictions. For example, our ENS^2^ showed around 34%/36% higher in Corr and 18%/15% lower in ER than MLspike/OASIS. When compared to CASCADE, our systems also showed superior performance for both spike-rate prediction and spike-event prediction (Figure 3A-D & Supplementary Figure 1C-D, p<0.0001 for all cases). In particular, our ENS^2^ showed around 5% higher in Corr and 4% lower in ER than CASCADE.

We took a deeper look into these results by considering the excitatory neurons and inhibitory neurons separately (Supplementary Figure 3). Generally speaking, all the four system performed worse in inhibitory neurons than in excitatory neurons (see Discussion). Notwithstanding, the ENS^2^ system presented better results in inhibitory neurons than the other models (Corr: p=0.013 for ENS^2^ vs. CASCADE, p<0.0001 for the rest; vRD: p=0.0093/0.0018 for ENS^2^ vs. CASCADE/OASIS, p<0.0001 for the rest; ER: p<0.0001 for ENS^2^ vs. MLspike; VPD: p<0.0001 for ENS^2^ vs. MLspike/OASIS). For example, our ENS^2^ showed around 11%/54%/27% higher in Corr and 6%/40%/2% lower in ER than CASCADE/MLspike/OASIS for inhibitory neurons. These results proved that our ENS^2^ system yielded better performance consistently for both excitatory and inhibitory neurons.

We also investigated how the performance of each algorithm varied for each single neuron (or dataset) (Figure 3E-H). The comparison showed that for neurons that were well/badly predicted by one algorithm were usually predicted (relatively) well/badly by the other algorithms as well (e.g. better on dataset 9-13 but worse on dataset 14-15). This suggests that the performance of inference for certain neuron (or dataset), regardless of algorithm, would depend significantly on their own properties. We explored the underlying factors to such phenomenon below (Figure 8-9).

We selected several specific segments of recording from five different neurons (Figure 2E-F, Figure 4) for further comparison among the four algorithms. In Figure 2E, the recording with GCaMP6s indicator has relatively high frame rate of ∼158Hz, and the original calcium trace (in *ΔF*/*F*_*0*_) has large amplitudes with only small noise. Figure 2F shows that CASCADE tended to output broader spike-rate prediction and thus broad spike-events than ENS^2^. This may be because the “noise matching input” used by CASCADE could contain more noise than the actual input due to rounding of noise level to integer values in the algorithm. As such, the model perceived more noise than actually existing in the inputs. On the other hand, both MLspike and OASIS significantly over-estimated with long sequences of spike-events. In contrast, our system (ENS^2^) showed better predictions than these three methods for all evaluation metrics (values on the right in Figure 2F). Figure 4A shows another sample recording of high SNR but with GCaMP6f indicator. In this case, our ENS^2^ system again recovered the spike-rate and spike-event characteristics the most properly, while the other three systems tended to under-estimate and missed several spikes. On the other hand, when the frame rate of the recording was decreased to 30Hz with lower SNR (Figure 4B), the inference task became more challenging. We observed slight over-estimation from the ENS^2^ system, and several missed spikes from the CASCADE system. However, both MLspike and OASIS performed poorly in this case, probably due to biased parameters during auto-calibrations. Figure 4C demonstrates an example with synthetic dyes and much lower frame rate at 7.8Hz, where the calcium dynamics are slow, and the signal quality reduces. This slow dynamic caused shifted predictions in time on ENS^2^, CASCADE, and MLspike. The low SNR caused MLspike to over-estimate, while OASIS failed to detect any spike in this segment. Nevertheless, the ENS^2^ system still recovered the firing pattern the best among the four. On the other hand, we have shown that these systems performed worse on inhibitory neurons (Supplementary Figure 3), where the firing rate is generally much higher with bursting. Figure 4D shows the inference results on an inhibitory neuron. Considering the low frame rate and slow dynamic of calcium imaging, the ground truth spiking activities would be extremely challenging to be recovered. Nevertheless, predictions from ENS^2^ and CASCADE resembled the temporal patterns of the ground truth much better than MLspike and OASIS, in that MLspike tended to under-estimate and OASIS tended to over-estimate. Together, we showed that our proposed ENS^2^ systems could maintain robust inference capability under varying conditions.

**Figure 4.**
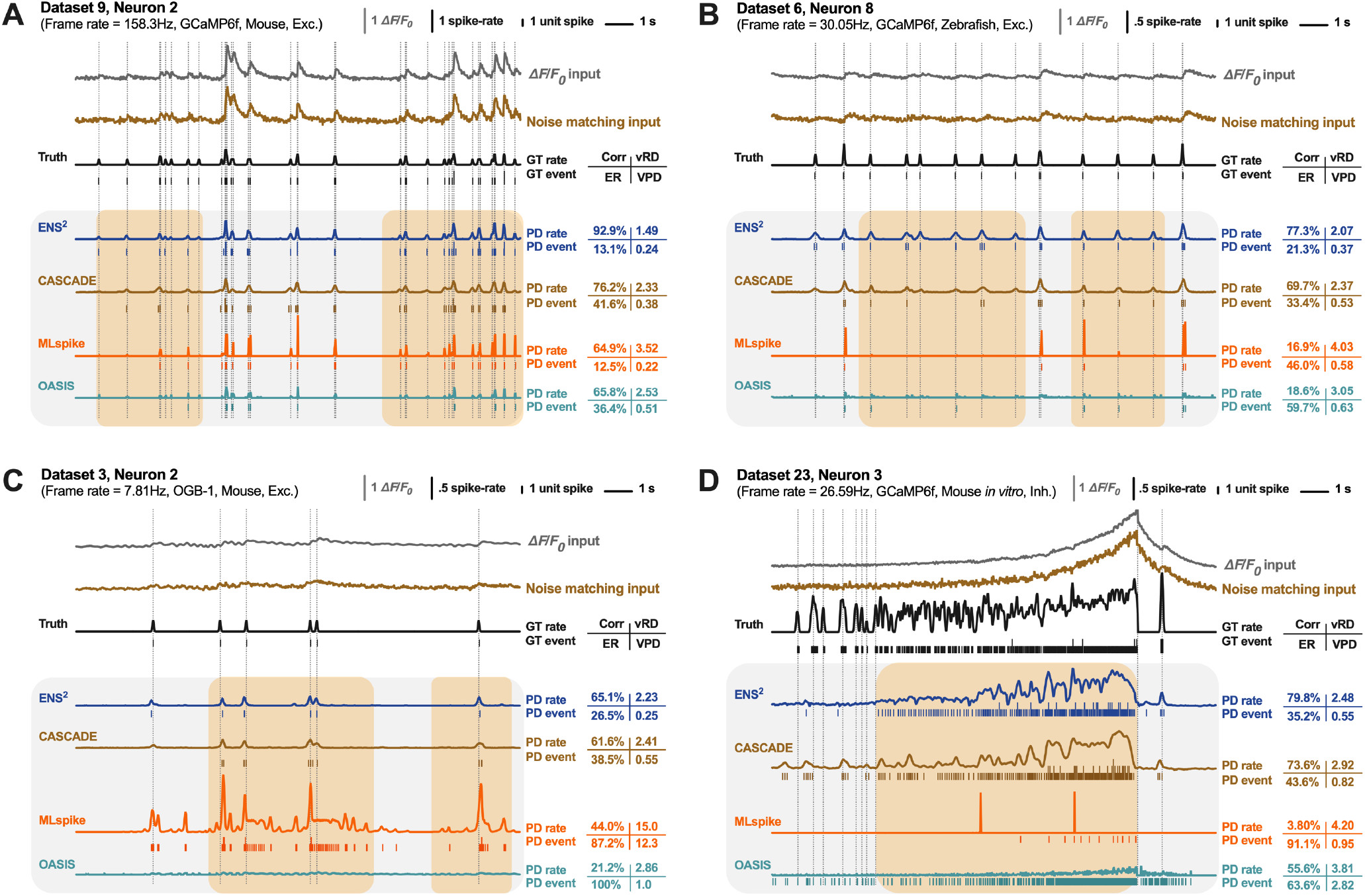
Examples of spike-rates and spike-events prediction from our proposed ENS^2^ and state-of-the-art methods. In each subfigure, *ΔF*/*F*_*0*_ calcium inputs and noise matching inputs are shown with the ground truth (GT) on top. The predicted (PD) spike-rates and corresponding spike-events by various methods are shown below. Metrics on the right measure the performance on the corresponding neurons. Orange shaded areas represent regions of interest where discrepancies in predictions are significant among different methods.

We would also like to point out that although our proposed ENS^2^ is data-driven, it is less computationally demanding than the previous method (e.g. CASCADE, Figure 5A). For a specific sampling rate (e.g. 60Hz), series of noise matching models were trained in CASCADE to meet the need of different noise levels. Each of their noise matching model consisted of 5 identical networks for ensemble learning to boost performance. On the other hand, only a single network is required in our ENS^2^ to predict data under each sampling rate for various conditions. As a result, the ENS^2^ with U-Net requires 20k (around 18%) fewer trainable parameters, and only a maximum of 5.12 million data segments are fed for training, two orders less than CASCADE (Figure 5A). In fact, the ENS^2^ system could perform at similar level with only 50k trainable parameters (around 30% of that in CASCADE) (Supplementary Figure 2). In particular, training of ENS^2^ on millions of samples during benchmarking was completed in around 2 minutes on average for each testing dataset, which was one-order faster than CASCADE (Figure 5B). This will enable cost-effective re-training or fine-tuning when more paired datasets are available to improve our model further.

**Figure 5.**
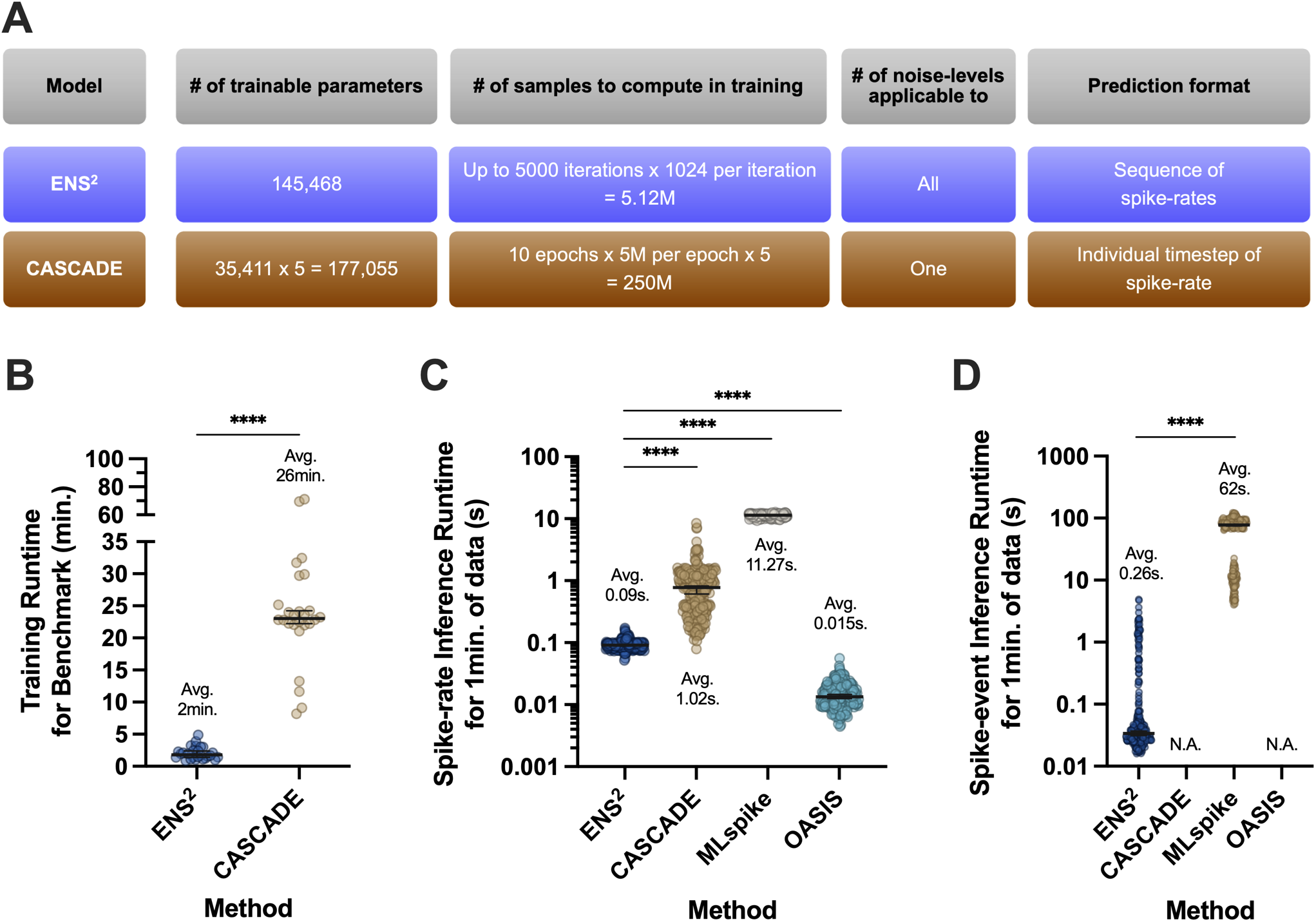
Computational complexity and runtime of ENS^2^ and the state-of-art algorithm. (**A**) Comparison of neural networks adopted in ENS^2^ and CASCADE. Our method requires two-order fewer samples to compute, and could generalize to all noise-levels. (**B**) Comparison of the runtime for training neural network models in ENS^2^ and CASCADE. (**C**) Comparison of runtime spent for spike-rate inference among different algorithms. The runtime is measured for each neuron and normalized to 1 minute. (**D**) Same as (**C**), but for spike-event inference. Colored circles present each neuron from all 26 datasets. Error bars present medians with 95% confidence intervals. Text denotes mean values of all neurons. Asterisks show the results of Friedman test with Dunn’s multiple comparisons (**C**) or two-sided Wilcoxon signed-rank test (**B & D**) between the indicated systems. The runtime was measured on a PC with an Intel Xeon E5 1630 v4 CPU and Nvidia GTX 1080 GPU.

We also evaluated the efficiency of these systems during inference (Figure 5C-D). For spike-rate inference, our ENS^2^ system took one-order less time than CASCADE to complete for every 1 min. of recording, and two-order less time than MLspike on average. The OASIS system showed the fastest spike-rate inference, despite compromised performance shown in our comparisons (Figure 2A-B, Supplementary Figure1 C-D). For spike-event inference, our ENS^2^ system with the greedy estimation algorithm (see Methods, Figure 1B) took two-order less time to complete than MLspike. The runtime was not measured for CASCADE as its built-in estimation algorithm took days to complete the benchmark for all datasets. The runtime was not presented for OASIS neither as the spike-event predictions were obtained by hard thresholding. Overall, we show that our ENS^2^ demonstrated better performance as well as computational efficiency for inferring spikes from calcium data than the state-of-art methods.

### Application to information encoding in primary visual cortex

In the above benchmark, we showed that ENS^2^ accomplished relatively high performance and high efficiency for inference in un-seen recordings. Next, we ask if our system would provide additional insights to physiological observations *in vivo* as previous models did (de Vries *et al*., 2020; Rikhye and Sur, 2015; Wilson *et al*., 2012). Here, we trained our full ENS^2^ system with all 20 excitatory datasets available, and then deployed it to unseen calcium imaging data recorded from the primary visual cortex (V1) (Figure 6-7 & Supplementary Figure 4). We collected *in vivo* calcium fluorescence images with GCaMP6s indicators from V1 of mice that were shown to drifting grating stimuli of four unique orientations, that move in two opposing directions (8 directions total) (Figure 6A, see Methods). Responsive neurons were selected, and their fluorescence signals were processed into calcium traces (*ΔF*/*F*_*0*_) for further analyses (Figure 6B1-2 & 6C1-2, see Methods). We then used our ENS^2^ system to predict the spike-rate (Figure 6B3 & 6C3) and spike-event (Figure 6B4 & 6C4) accordingly. We compared our analyses with these different inputs and verified if the spike inference has any positive impact in understanding the information encoding in V1 than the raw *ΔF*/*F*_*0*_ trace alone. Here, two presentative neurons are shown in Figure 6B-C.

**Figure 6.**
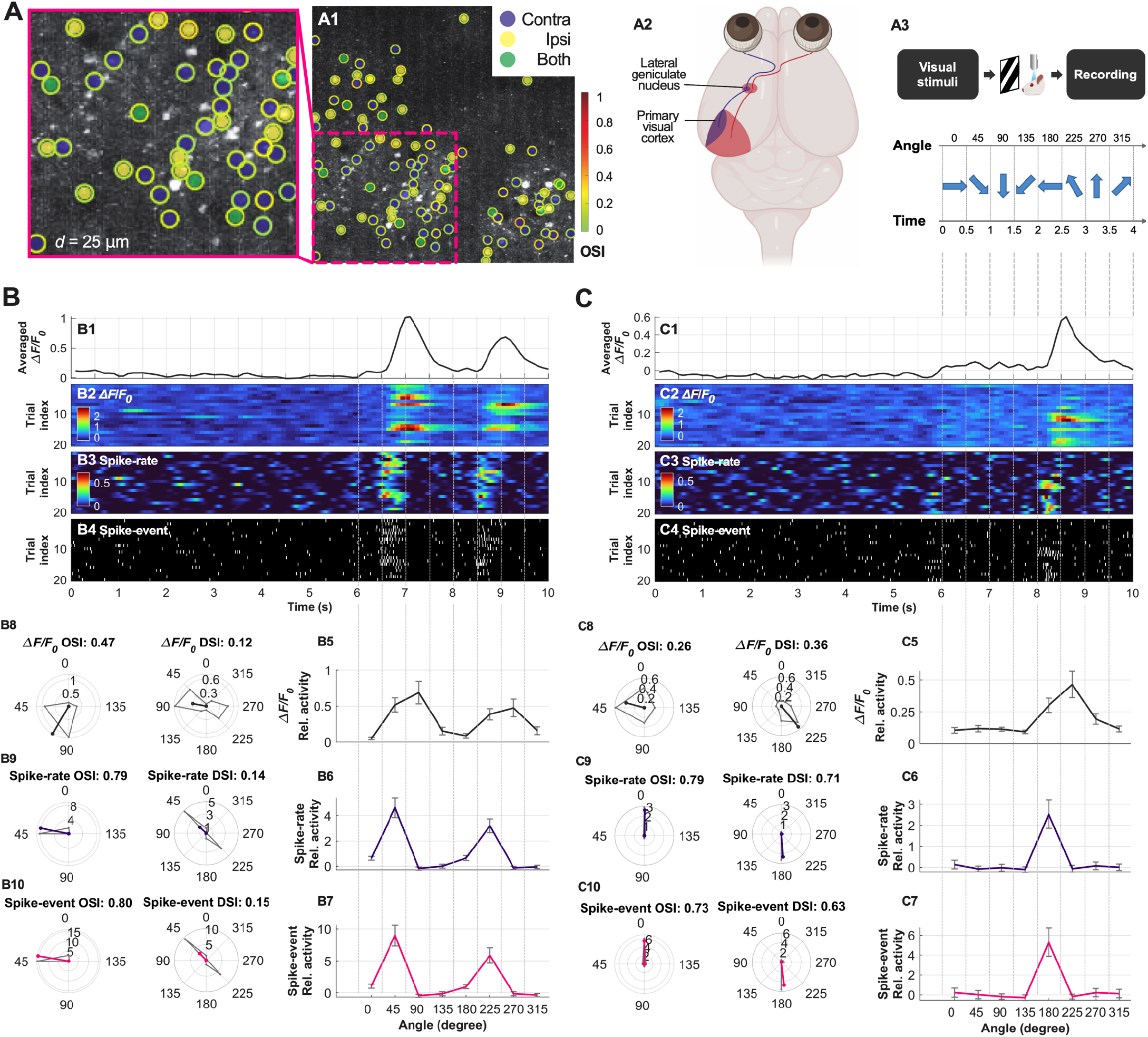
Spike inference with ENS^2^ for calcium imaging data collected in primary visual cortex (V1) in a visual stimulating experiment. (**A**) An example of image (**A1**) recorded from the binocular zone of the left V1 (**A2**) of mice subject to visual grating stimuli (**A3**). Visually responsive neurons are labeled in (**A1**) and are considered for further analyses (see Methods). The inner color denotes the response type of the neurons. *Contra* and *ipsi* refers to the neurons that are responsive to contralateral (right) and ipsilateral (left) eye inputs, respectively. *Both* means the neurons are responsive to both sides of inputs. The outer color denotes the OSI computed from *ΔF*/*F*_*0*_ signal for that neuron. (**B**-**C**) Examples of recorded calcium signals and the predicted spike-rates and spike-events by ENS^2^ (**B1**-**B4, C1**-**C4**). Tuning curves and selectivity indexes (**B5**-**B10, C5**-**C10**) are computed based on three types of inputs (**B2**-**B4, C2**-**C4**), respectively.

**Figure 7.**
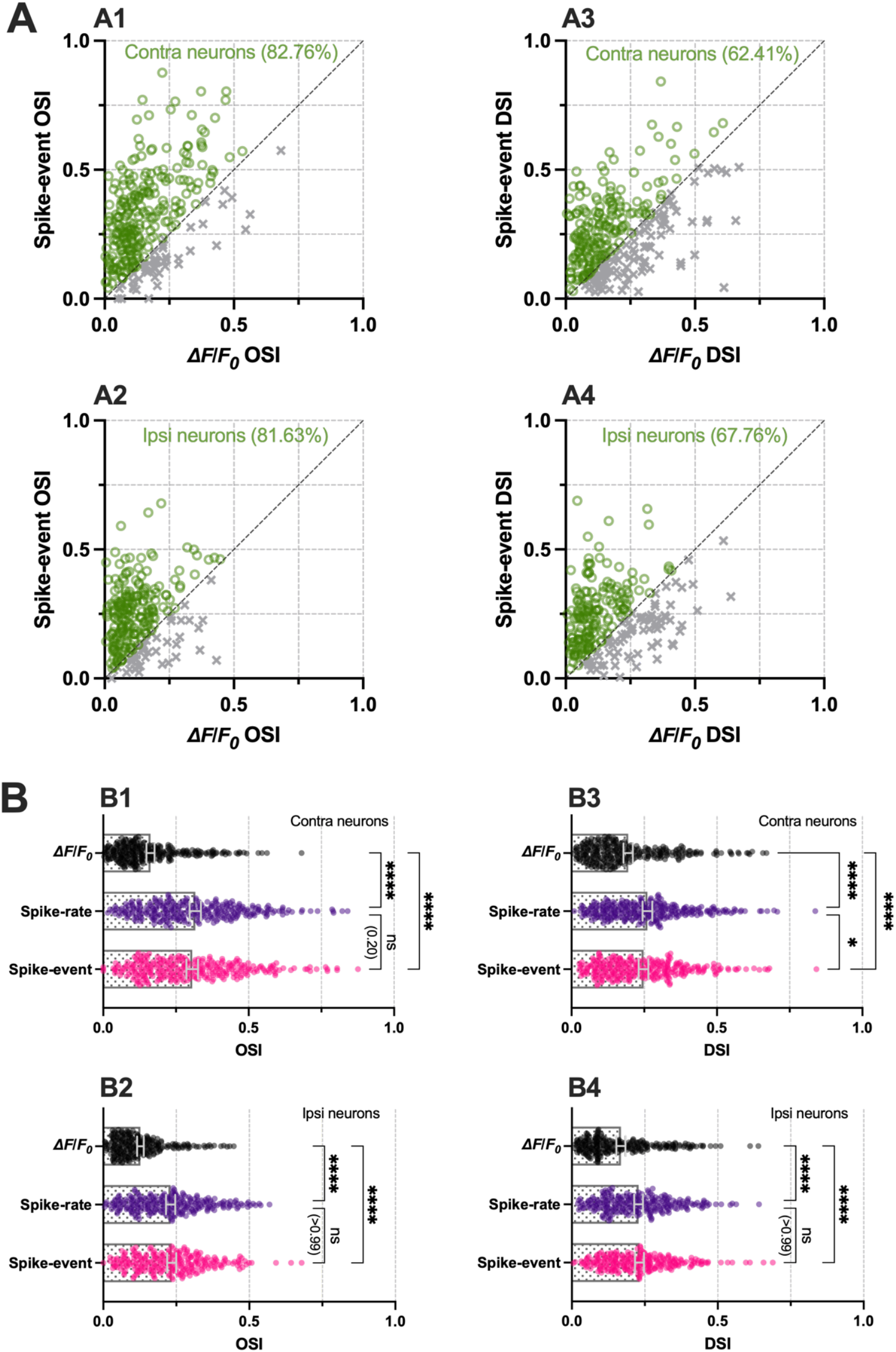
Quantitative analyses of the inferred neurons with ENS^2^ in the visual stimulating experiment. (**A-B**) Comparisons of OSI/DSI computed from *ΔF*/*F*_*0*_ signal and after spike inference with ENS^2^ for all the 290 *contra* and 245 *ipsi* neurons considered. Each circle/cross/dot represent one recorded neuron. Asterisks show the results of Friedman test with Dunn’s multiple comparisons between the indicated predictions.

We first constructed the tuning curves (see Methods) for each neuron (Figure 6B5-7 & 6C5-7). It was observed that the resultant tuning curves were broader when computed using *ΔF*/*F*_*0*_ (Figure 6B5 & 6C5) than using spike-rate or spike-event (Figure 6B6-7 & 6C6-7). In particular, the spike-rate/spike-event tuning curves exhibited sharpened preferred orientation for these cells. The broader tuning curve by *ΔF*/*F*_*0*_ was mainly due to the long “tail” (decaying edges) in *ΔF*/*F*_*0*_ signal after each peak (Figure 6B1 & 6C1) resulting from the slow dynamics of calcium indicators. Consequently, the long “tail” of *ΔF*/*F*_*0*_ may lead to a shift in the preferred orientation and broader tuning curves. On the other hand, by predicting the spike-rate/spike-event with our ENS^2^ system, we successfully removed these long tails in the signal. We further quantified the preferred orientations by computing the orientation selectivity index (OSI) and direction selectivity index (DSI) from the tuning curves for each neuron (see Methods). The neuron in Figure 6B is a sample cell exhibited high OSI but low DSI; while the neuron in Figure 6C is exhibiting high OSI and high DSI (Mazurek *et al*., 2014) (see Methods & Supplementary Figure 7). Our results show that OSI computed from *ΔF*/*F*_*0*_ was lower for both neurons in Figure 6B-C but was higher when computed from spike-rate/spike event (from 0.47 to 0.79/0.80 and from 0.26 to 0.79/0.73, respectively). Similarly, DSI computed from *ΔF*/*F*_*0*_ was lower for the neuron in Figure 6C, but increased when computed from spike-rate/spike event (from 0.36 to 0.71/0.63). Supplementary Figure 4A-B showed two neurons that appeared to have a preferred orientation at 0°, but with a strong decaying tail in their *ΔF*/*F*_*0*_. Due to such up-shifted amplitude in *ΔF*/*F*_*0*_, the computed OSI or DSI tends to be higher. In contrast, the predicted spikes removed the up-shifting, and resulted in a more clear-cut readout of the OSI/DSI measurement (such as Supplementary Figure 4B).

This observation is consistent for the neuron population we have recorded (290 *contra* and 245 *ipsi* neurons (see Methods)). Figure 7A1-2 showed that OSI computed with spike-event was higher than that computed with *ΔF*/*F*_*0*_ in >80% of the cells (82.76% for *contra* neurons and 81.63% for *ipsi* neurons). Also, Figure 7A3-4 show that DSI computed with spike-event was higher than that computed with *ΔF*/*F*_*0*_ in >60% of the cells (62.41% for *contra* neurons and 67.76% for *ipsi* neurons). Overall, the mean OSI/DSI for both *contra* and *ipsi* neurons were higher with spike-event/spike-rate than *ΔF*/*F*_*0*_ (Figure 7B, p<0.0001 for all cases). From the results of all these cases, we believe that the OSI and DSI computed from spike-rate/spike event predicted from our ENS^2^ system would discriminate the response pattern of these neurons better than those derived from the original *ΔF*/*F*_*0*_.

It is also worth noting how the spike inference would benefit analyses of neurons that show weak responses or where the signal-to-noise ratio (SNR) is low (Supplementary Figure 4C-D). In Supplementary Figure 4C, the *ΔF*/*F*_*0*_ of this neuron only varied over a range of 0.2 such that the SNR was very low. The resultant noisy tuning curves suggested imprecise orientation or direction selectivity for this neuron. Instead, the spike inference increased the response SNR such that the tuning curve was much sharpened, showing preferred orientation at 0° and 180° and hence an increased OSI. On the contrary, in Supplementary Figure 4D, the tuning curve from *ΔF*/*F*_*0*_ resulted in a sizable OSI (0.43) which may (falsely) suggest that this neuron has an orientation preference at 0°. Nevertheless, after spike inference with our ENS^2^, the small peaks in the original *ΔF*/*F*_*0*_ signal were filtered out, resulting in negligible OSI and DSI. This suggested that this neuron in fact has very weak selectivity and may not be considered a real responsive cell. In this sense, the spike inference by our ENS^2^ increased the SNR of the neuronal response to not only improving the sensitivity in detecting the orientation selectivity of the neurons, but also to screen out some marginally responsive neurons.

### Factors affecting inference performance

To understand further what contributed to good spike inference performance for data-driven methods, we investigated how the data itself affected the performance of these models. These insights may further facilitate data collection and preparation for improving data-driven models (e.g. our ENS^2^).

Figure 8A-D show the performance with individual dataset achieved by different configurations of models (the configurations are numbered horizontally in the same order as in Figure 2A-D). The results show that performance indeed depended strongly on the dataset. For instance, regardless of the networks used, dataset 15 and 17 achieved notably worse vRD than other datasets (Figure 8B), and some datasets (e.g., 21-22) showed considerably worse ER than the others (Figure 8C). These were also observed when comparing our ENS^2^ to state-of-the-art methods (Figure 3E-H). We extracted a number of parameters from the calcium recording for each neuron, such as noise level, peak firing rate, frame rate, and calcium transient amplitude (see Methods), and examined how they may affect the inference performance (Figure 8E-H, Supplementary Figure 1I). It is shown that the noise-level and peak firing rate had less influences on the performance. On the contrary, the frame rate correlated with the inference performance more significantly, probably because higher frame rate provides higher SNR. Importantly, we found that the transient amplitude of the dataset was the key predictor for the inference performance (including Corr, vRD, ER, and VPD). We also noticed that the original transient amplitudes in *ΔF*/*F*_*0*_ were of critical importance for inference, which related the number of spike events for a certain calcium indicator. A previous model-based study also testified that an accurate estimate of transient amplitude improved performance (Éltes *et al*., 2019). The transient amplitude indeed strongly depends on the calcium indicators’ sensitivity. It is clear that one major bottleneck of inference algorithm is in the calcium indicators.

We next investigated the preferred ways of supplying the training data for our system. We first examined this by training the model using three different subsets of the training data for each of the testing data, according to the types of calcium indicators. Here, “*All*” used all the possible training set in training (same as the default leave-one-dataset-out protocol) while “*Same*” used only those training datasets with the same calcium indicator as the testing data. Note that these simulations were performed separately for datasets with excitatory and inhibitory neurons, respectively (see Methods). Also note that some of the indicator comprise only one dataset (e.g. GCaMP5k) and hence they were not tested in the “*Same*” protocol. Figure 9B-D and Supplementary Figure 1J show that “*Same*” did not perform better than “*All*” significantly in general. Similar observation was found in (Rupprecht *et al*., 2021) where they reported that clustering the same calcium indicators for training showed no advantage in CASCADE. It seems that a generalized inference model (“*All*”) is sufficiently good for the inference task than using multiple indicator specific models (“*Same*”). The advantage of indicator specific training was most prominent in dataset with GCaMP6f indicators (Figure 9A-E, Supplementary Figure 1J-K). This was probably because GCaMP6f is the dominating indicator in the whole benchmark database. Nevertheless, the difference remained quite small. As such, we recommend training the neural network based model (e.g. our ENS^2^ system) with all available paired data to exploit their generalization capability.

**Figure 8.**
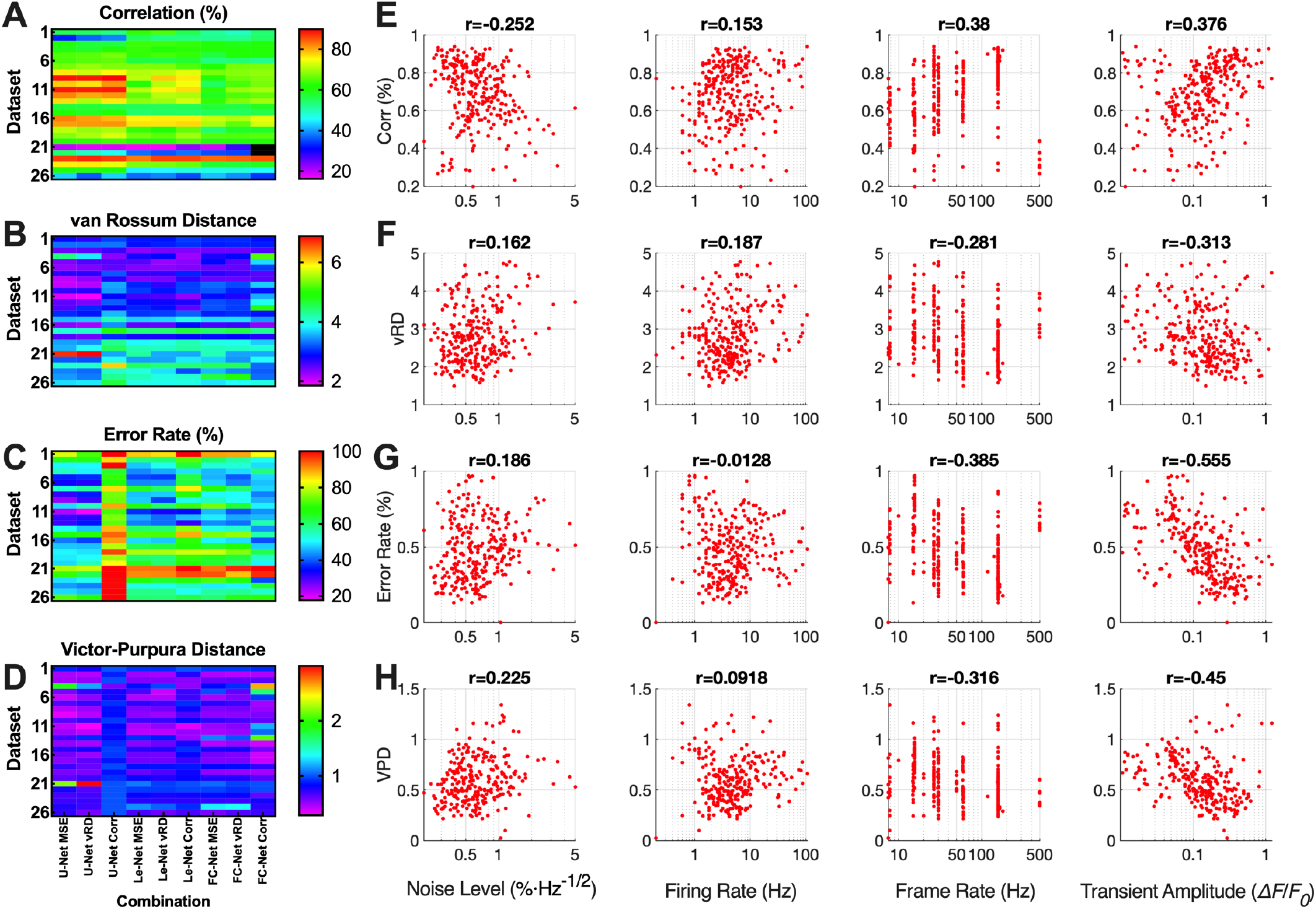
Effect of dataset properties on spike inference performance. (**A-D**) Performance with different configurations (see **Figure 2A-D**) (x-axis) on each dataset (y-axis). Colormap shows the performance of the median neurons of each dataset. (**E-H**) Spearman rank correlation coefficients (r) between the four properties of each dataset and the corresponding spike inference performances. Red dots represent each neuron from all datasets.

**Figure 9.**
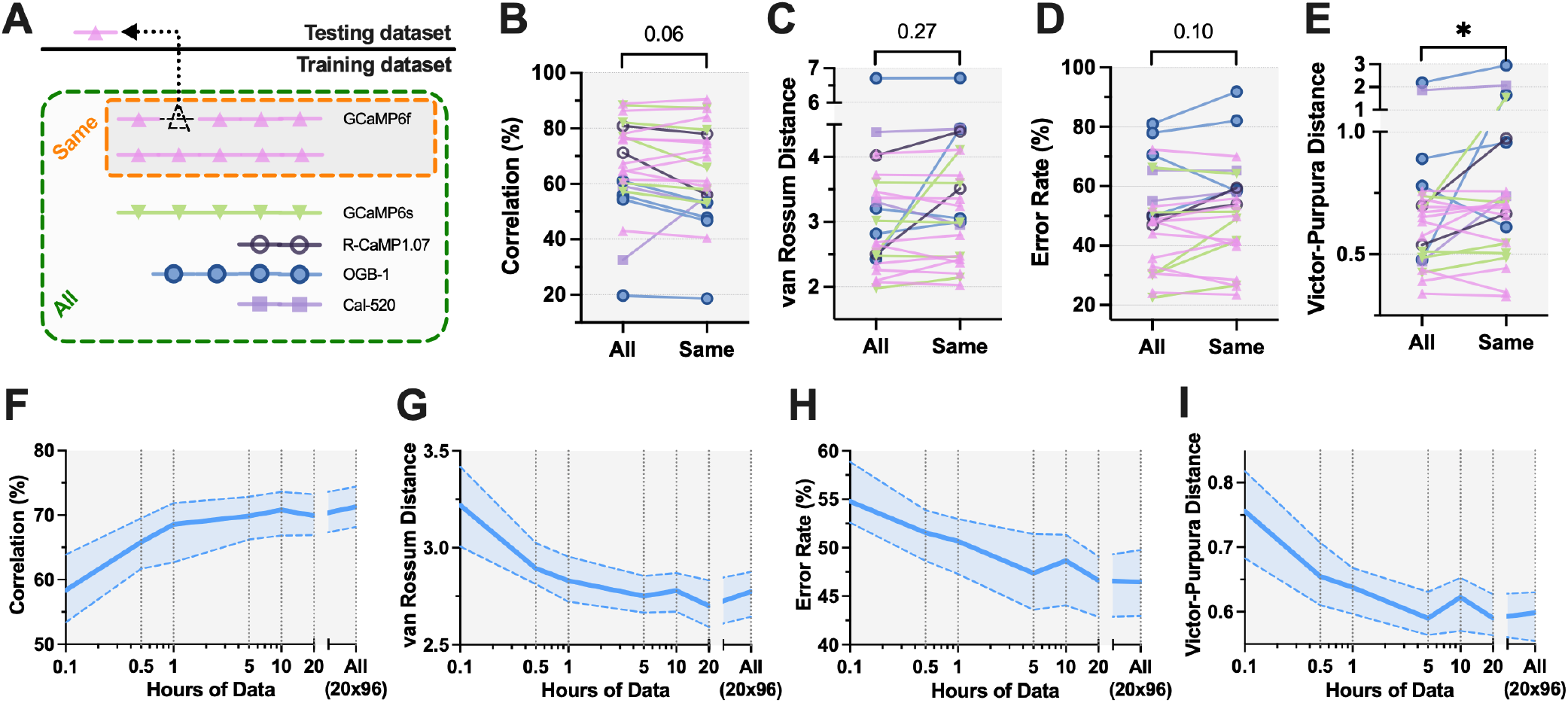
Effect of calcium indicators and data quantity on spike inference performance. (**A**) An example to illustrate how the division of datasets was made based on calcium indicator when a GCaMP6f dataset was regarded as testing dataset. When testing on an excitatory/inhibitory dataset, *All* refers to all the other 19/5 datasets, *Same* refers to the datasets that also used GCaMP6f. Note that the simulations were performed separately for excitatory/inhibitory datasets. (**B-E**) Performance of spike inference for different types of calcium indicators. Colored circles present the performance of median neurons of each testing dataset. Asterisks show the results of two-sided Wilcoxon signed-rank test between the indicated partitions. (**F**-**I**) Performance of spike inference with different length of training data. Shaded areas denote medians with 95% confidence intervals.

Since our ENS^2^ is a data-driven model, we also wondered how much training data is needed for achieving good inference performance. We randomly sampled different numbers of segments from the total of over 20 hours of available paired data from all excitatory datasets. When supplying all available paired data to the model, the total duration was approximately 20 × 96 hours since the paired data was segmented with a step of 1 data point (see Method). Not surprisingly, the performance of inference increases with the amount of training data, but it converges with roughly 5 hours of paired data (Figure 9F-I, Supplementary Figure 1L-M).

Several other factors may also have significant impact on the inference performance, such as sampling rate (resolution of prediction), size of smoothing window (for spike-rate prediction), and hyper-parameter of evaluation metric (e.g. ER window size). The comparisons are summarized in Figure 10A-D. Figure 10A shows that the Corr increased consistently with larger smoothing windows. Similar observations can also be found in several recent studies (Berens *et al*., 2018; Stringer and Pachitariu, 2019; Theis *et al*., 2016). This is because the GT spike-rates convolved from the GT spike-events with larger smoothing windows have smoother and broader patterns, which favored the measure of Corr. Figure 10F shows the GT spike-rates obtained by convolving the GT spike-events in Figure 10E with varying smoothing window sizes (25ms to 200ms) and their corresponding predictions. Apparently, the smoother and broader waveform of GT spike-rate (with larger smoothing windows) simplified the prediction task, and it was easier to achieve a high Corr with such simpler and smoother PD spike-rate waveform. This was also true for the spike-rate evaluation with vRD and Error (Figure 10B & Supplementary Figure 1N).

**Figure 10.**
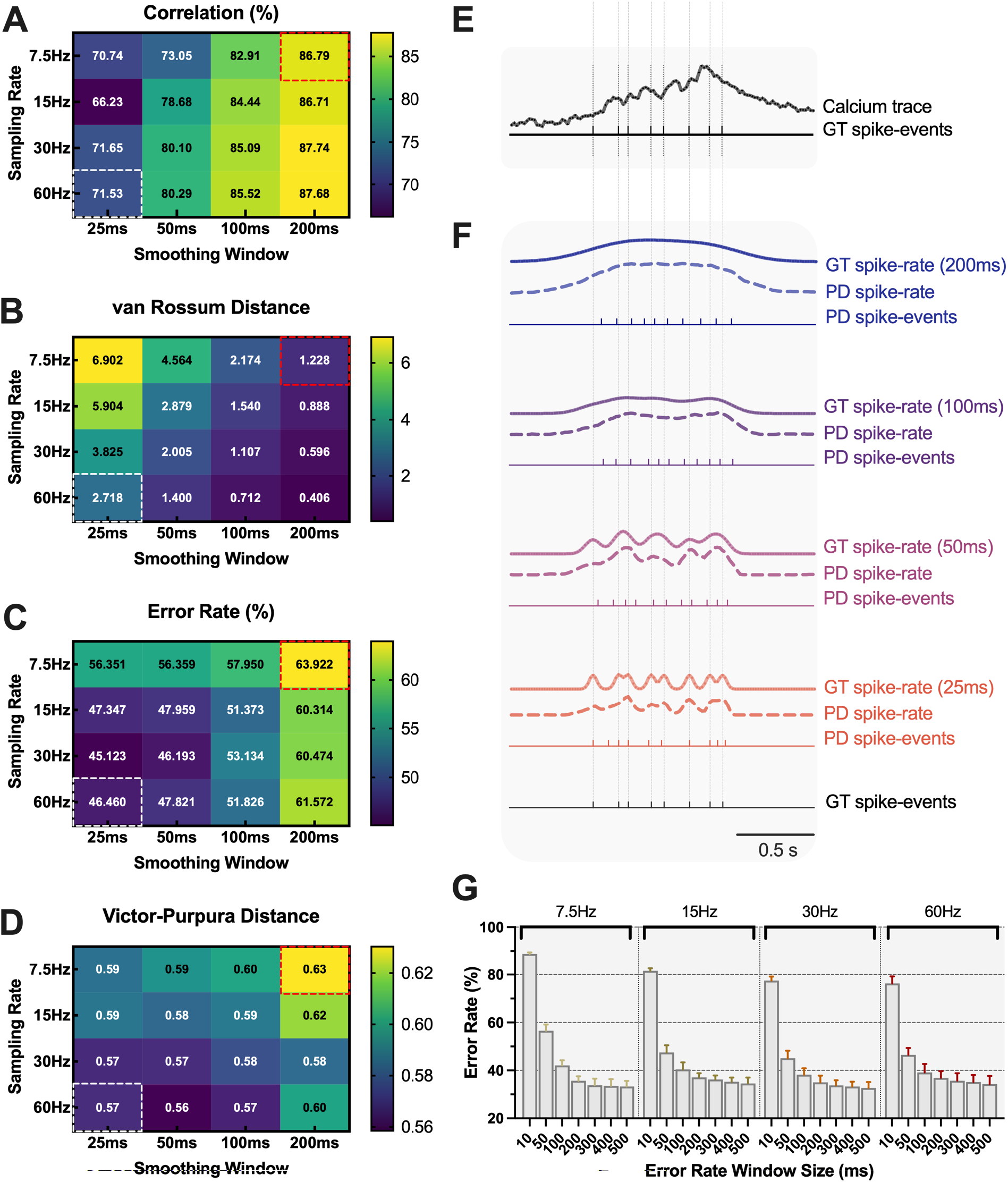
Effect of evaluation metrics on spike inference performance. (**A-D**) Performance of correlation, van Rossum distance, error rate, and Victor-Purpura distance, respectively, when measured under different sampling rates and smoothing window sizes. Embedded values show median performance among all neurons. White and red squares denote the evaluation schemes adopted by ENS^2^ and CASCADE, respectively. (**E**) Example of calcium signals under 60Hz with paired spike-events. (**F**) Examples of ground truth spike-rates convolved with different smoothing window sizes (from 200ms to 25ms). The resultant spike-rate and spike-event predictions are also shown. (**G**) Performance of error rate when measured under different sampling rates and error rate window sizes.

However, we argued that such resultant “better” performance (e.g. high Corr) would not always guarantee meaningful predictions as reflected in the PD spike-events, since multiple GT spike-events could be merged into a single peak of spike-rate (Figure 10E-F). Instead, the temporal firing patterns could be better predicted with narrower smoothing windows. On the other hand, spike-event predictions (VPD & ER, Figure 10C-D & Figure 10G) generally improved with higher sampling rates. This is quite reasonable as smaller bin sizes allow more precise estimation of spike-events from spike-rate predictions. Moreover, when high sampling rates were used (e.g. 30 or 60Hz), VPD and ER also reduced along with smoothing window sizes, indicating improved spike-event predictions. Here, the spike-event inference performance would possibly be restricted by the overly smoothed spike-rates (e.g. Figure 10F). We also analyzed the effect of ER window sizes on ER evaluation (Figure 10G). As expected, smaller ER window sizes put higher demand on the evaluation of the algorithm and hence, resulted in larger ER, which is similar to the findings reported in a previous study (Deneux *et al*., 2016).

Given these analyses, we suggest that our ENS^2^ system should be trained with inputs at sampling rate of 60Hz (by resampling when necessary) with 25ms smoothing windows for practical use (labeled with white dashed boxes in Figure 10A-D & Supplementary Figure 1N-O). Further increase in the sampling rate would cause computational over-head while to reduce the smoothing window size further might be harmful to training neural networks with gradient descent. We also repeated our benchmark using Causal smoothing kernels as in CASCADE (Rupprecht *et al*., 2021). Nevertheless, we found that using Gaussian smoothing kernels generally outperform Causal smoothing kernels (Supplementary Figure 5, p<0.001 for Corr/ER/Error; p<0.0001 for vRD). It is also worth noting that the CASCADE algorithm (Rupprecht *et al*., 2021) was indeed benchmarked under 7.5Hz with 200ms smoothing windows. Supplementary Figure 6 shows that our ENS^2^ consistently outperformed the CASCADE algorithm at these settings for both spike-rate and spike-event predictions (Corr: p=0.0058; vRD/Error/Bias: p<0.001; ER/VPD: p<0.0001) (also labeled in red dashed boxes in Figure 10A-D & Supplementary Figure 1N-O). These results support that our ENS^2^ is a versatile and highly effective algorithm for spike inference from calcium signals.

## Discussion

In this work, we have developed a high performance inference system (ENS^2^) through extensive and empirical research, and showed its usefulness in inferring both spike-rates and spike-events.

We have found that networks with convolutional layers (e.g. U-Net and Le-Net) typically out-performed the other (e.g. FC-Net). This may be partly due to the regularization capability of the convolutional layers. In other words, it provides larger receptive fields (with context information such as calcium dynamics) with fewer trainable parameters and constrained kernel shapes. On the other hand, it is quite intuitive for humans to examine the calcium segments fraction by fraction to identify spike-events, just as sliding a kernel for convolution by the artificial neural networks. In fact, recent data-driven models (CASCADE (Rupprecht *et al*., 2021), S2S (Sebastian *et al*., 2021)) also used a network with convolutional layers. We speculate that the state-of-art performance of ENS^2^ also benefits from the skip-connection structure and sequence-to-sequence prediction manner of our modified U-Net. Recently, a 3D U-Net based model has also been proposed to improve SNR in calcium images and facilitate calcium signal extraction (Li et al., 2020). On the other hand, we revealed in our results that MSE loss could readily regulate the optimization of such models. While Corr is indisputably a major evaluation metric for spike inference, we suggest that using Corr as the sole loss function for deep learning models (e.g. in S2S (Sebastian *et al*., 2021)) might be defective in real world tasks. For example, the inferred spike-rates are illy-scaled in amplitude and are unable to recover spike-events faithfully.

While inferring spikes from calcium signals is a typical sequence-to-sequence translation task, recurrent neural networks (RNN) (e.g., LSTM based models (Berens *et al*., 2018)) may be also applied here. Nevertheless, one major bottleneck is that RNN should iteratively compute on each data point along time. Thus, the computational efficiency would be quite low compared to CNN based models. Similar issue is also faced by the powerful Transformer based models (Vaswani et al., 2017) designed for sequence-to-sequence translation, which have overly large numbers of parameters for online application of spike inference. On the contrary, in this work, we used deep convolutional networks with sequence-to-sequence translation ability (e.g., 1D U-Net), and it showed state-of-art performance with favorable efficiency.

Inhibitory neurons typically have higher instantaneous firing rates (Table 1), which complicated the spike inference task. Furthermore, the low frame rate and slow dynamics of calcium imaging may cause losses of even more details of the high frequency spiking activities (lower SNR, Figure 4D). As a result, the algorithms could only recover spikes from a smaller portion of information. On the other hand, the high firing rates raise challenges to algorithms when discriminating the actual units of spikes in a single time-step (time bin), since the transient amplitudes may change non-linearly as the number of spikes increases. It was shown that to recover every single spike from the calcium trace would be extremely difficult (Figure 4D). However, systems (e.g. ENS2) that could properly predict the temporal firing patterns at a longer time-scale for inhibitory neurons are still valuable tools for neuroscience research. Future work could explore how to incorporate the biophysical properties of these bursting pattern into the neural network, which will potentially improve the inference performance in the inhibitory neurons (Rahmati et al., 2016).

For actual applications, data-driven methods (e.g. ENS^2^ and CASCADE) are off-the-shelve for use without the need of further calibration, thanks to the generalization ability of neural networks, although CASCADE would need pre-training multiple versions to cater different noise-levels. On the other hand, for model-based methods (e.g. MLspike and OASIS), it is often necessary to define the parameters empirically or calibrate using some pre-defined algorithm for fresh recording, where ground truth paired data is normally unavailable. The similar scenario was encountered in our leave-one-dataset-out benchmarking where the testing dataset was separated from the training set. Interestingly, we found both tested model-based systems tend to under-estimate the spike-rate and/or spike-events (Supplementary Figure 1D & 1F). The two data-driven systems appear to perform better on these unseen data, which partly demonstrate their better generalization capability in this application.

Importantly, we have demonstrated that our spike inference algorithm could improve the analyses of real world calcium data such as in the study of neuronal orientation preference in primary visual cortex (Figure 6-7 & Supplementary Figure 4). Our results demonstrate that our algorithm can help perform analyses with both high throughput (from calcium imaging) and high precision (from spiking activities) in the study of our brain.

Several potential improvements for ENS^2^ are outlined as follows. Since ENS^2^ is a data-driven model, its performance would fluctuate upon different training data with various quality, diversity, as well as pre-processing procedures, etc. The benchmark database (Table 1) used in this work is essential to produce promising inference performance with the ENS^2^ system, yet further improvement could also be made. We have resampled both the training data and testing data to 60Hz for good quality inference (Figure 10). The resampling was performed with a Fourier based method (see Methods), which may not provide notable additional information to our models. We suggest that adopting diffusion based probabilistic models designed for biomedical time-series signal forecasting and imputation (e.g. (Alcaraz and Strodthoff, 2022; Ho et al., 2020; Tashiro et al., 2021)) may strengthen such resampling processes with temporal dependency. On the other hand, the effect of synthetic training data on our models remains unknown. It is possible that including synthetic paired data (e.g. NAOMi (Charles et al., 2019)) and/or generative models (e.g. generative adversarial networks (Goodfellow et al., 2014)) for data augmentation may further improve the generalization capability of our inference systems. Lastly, our inference system is trained and to be used in an end-to-end translation manner, where the inputs are *ΔF*/*F*_*0*_ calcium traces (see Methods). In fact, extracting *ΔF*/*F*_*0*_ signals from calcium imaging is another non-trivial process, in addition to the spike inference task here. During this extraction process, the quality of the resultant *ΔF*/*F*_*0*_ calcium trace may limit and/or bias the inference of our system. We therefore envision that modifying our ENS^2^ to directly take inputs from the source image-end (e.g. (Pnevmatikakis, 2019)) and output to the spike-end, may allow further exploitation and utilization of extra and unbiased information for spike inference.

## Acknowledgement

This work was supported by Research Grants Council of Hong Kong SAR Project CityU 11104220 (C.T.), ECS CUHK 24117220 (J.I.), and City University of Hong Kong (Project 7005645), Lo Kwee-Seong Biomedical Research Fund (J.I.) and Faculty Innovation Awards (FIA2020/A/04) from the Faculty of Medicine, CUHK (J.I.). Data and source codes to replicate the primary results will be shared upon reasonable request.

## AUTHOR CONTRIBUTIONS

C.T. conceived the study. Z.Z. designed the algorithms and performed analyses. J.I. and K.T. collected the *in vivo* calcium imaging data. J.I., K.T. and H. M.Y. pre-processed the data for spike inference and further analysis. M.S. provided expertise and inputs on dissecting V1 physiology, and visual stimulation design. All the authors contributed to interpreting the results and writing the manuscript.

## DATA AVAILABILITY

All data and source code to produce the benchmark and ENS^2^ system are available on https://github.com/tinlab/ens2 and the linked repositories therein. We used a publicly available database on https://github.com/HelmchenLabSoftware/Cascade (Rupprecht *et al*., 2021).

## SUPPLEMENTARY FIGURES

**Supplementary Figure 1.**
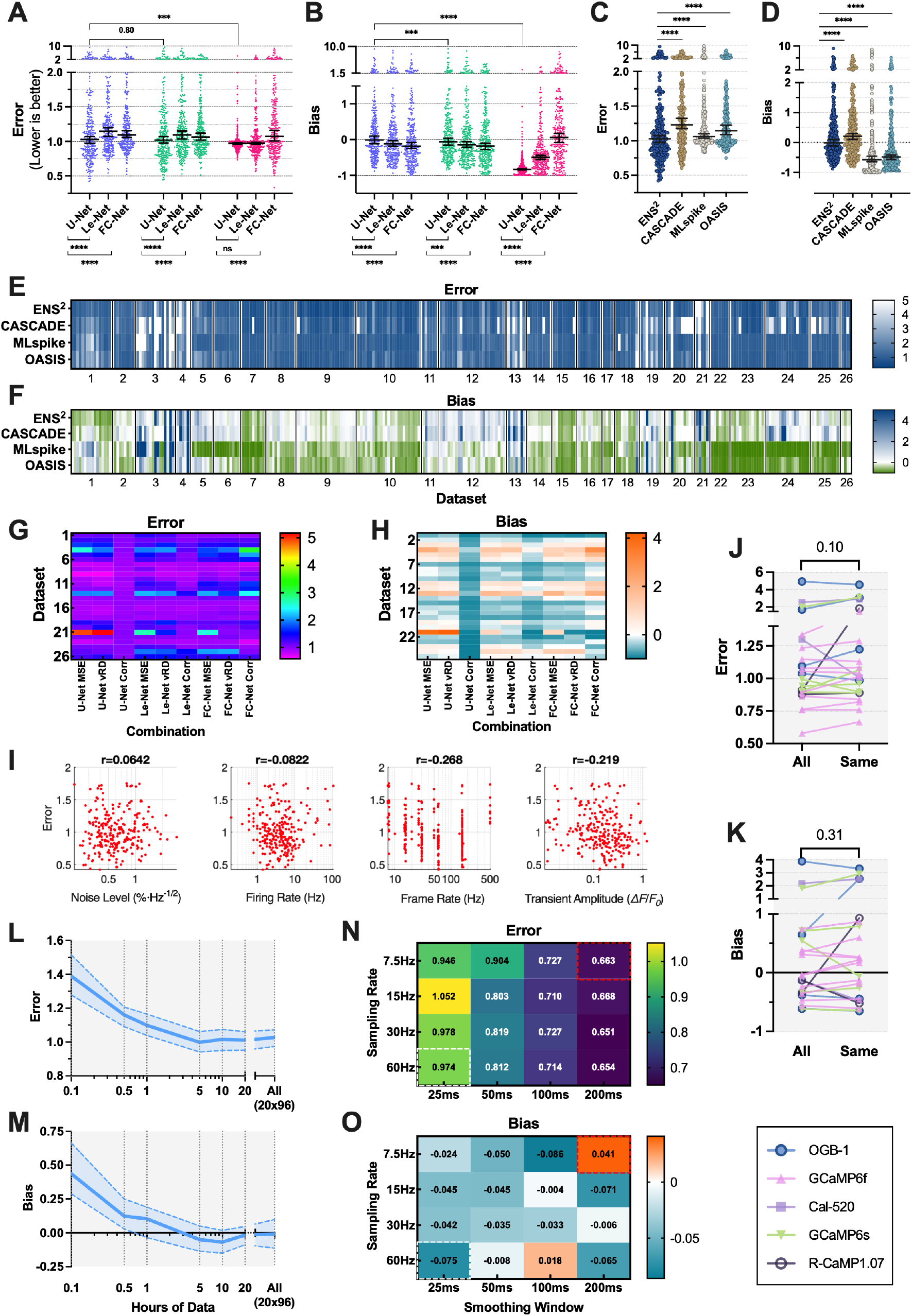
Results and analyses regarding Error and Bias assessment. Extension of (**A**-**B**) **Figure 2**, (**C**-**F**) **Figure 3**, (**G**-**I**) **Figure 8**, (**J**-**M**) **Figure 9**, (**N**-**O**) **Figure 10**, respectively. Conventions are same as in the corresponding figures.

**Supplementary Figure 2.**
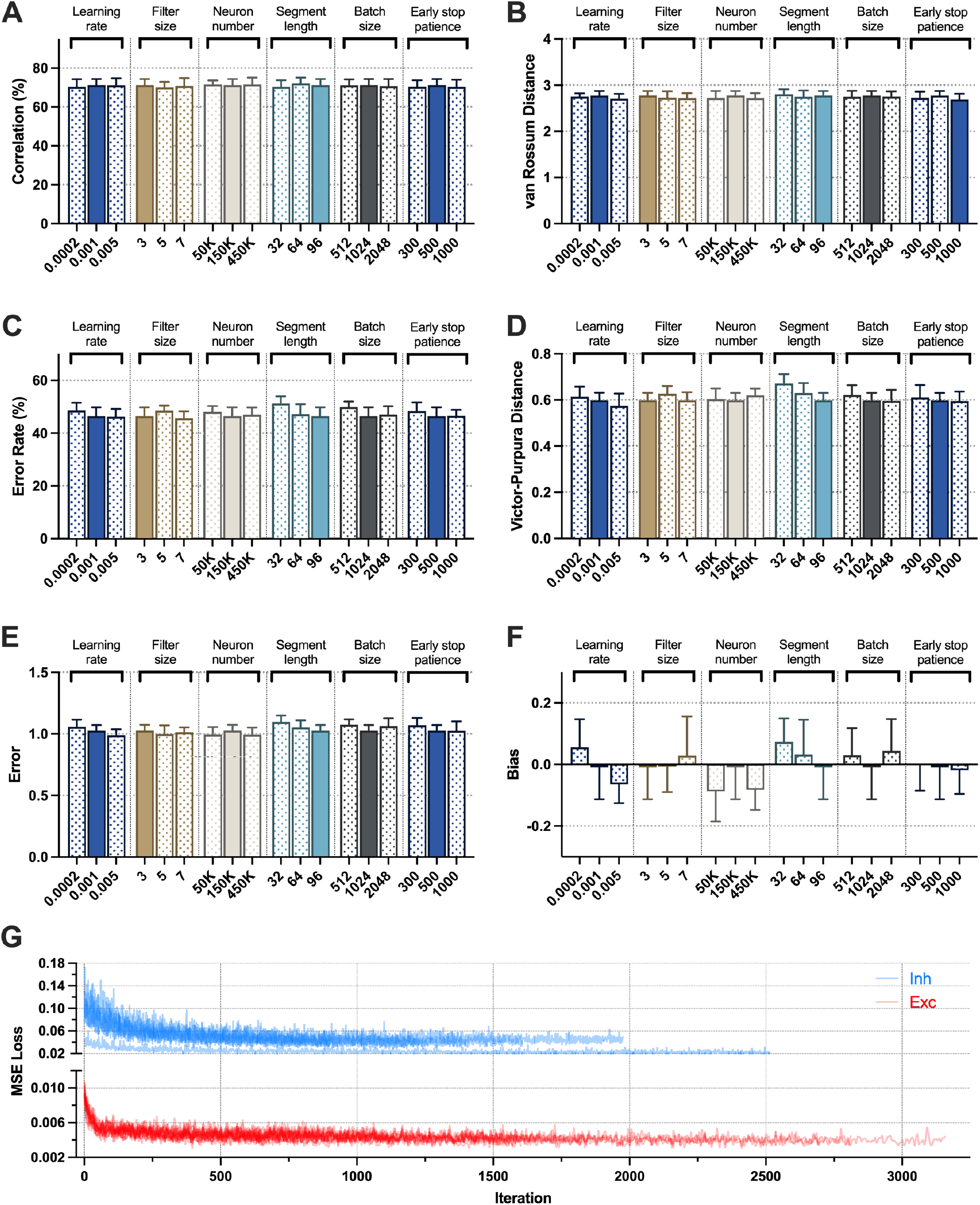
Performance comparison of the proposed method using various hyper-parameters. (**A-F**) Summary view of performance measured in correlation, van Rossum distance, error rate, Victor-Purpura distance, error, and bias, respectively. Results are obtained by ENS^2^. Filled bars represent default parameter values adopted in this study. (**G**) Dynamic loss curves for all datasets when training models with ENS^2^. Bar plots present medians with 95% confidence intervals.

**Supplementary Figure 3.**
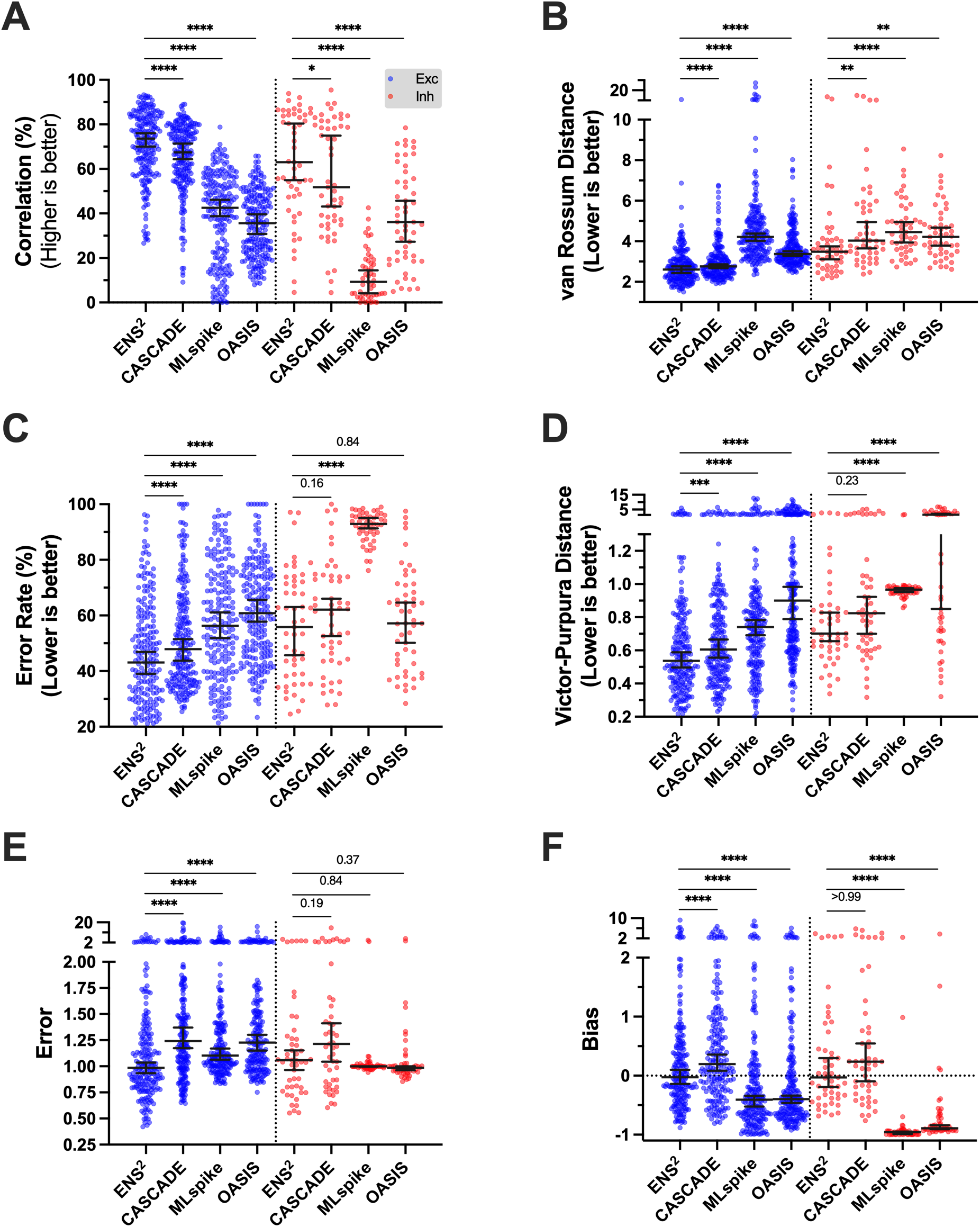
Performance comparison of our proposed ENS^2^ system against state-of-the-art algorithms when inferencing on excitatory and inhibitory neurons. Inference performance for excitatory (Exc) and inhibitory (Inh) neurons measured in (**A**) correlation, (**B**) van Rossum distance, (**C**) error rate, (**D**) Victor-Purpura distance, (**E**) error, and (**F**) bias, respectively. Colored dots denote the performance for each individual neuron. Error bars present medians with 95% confidence intervals. Asterisks show the results of Friedman test with Dunn’s multiple comparisons between the indicated systems.

**Supplementary Figure 4.**
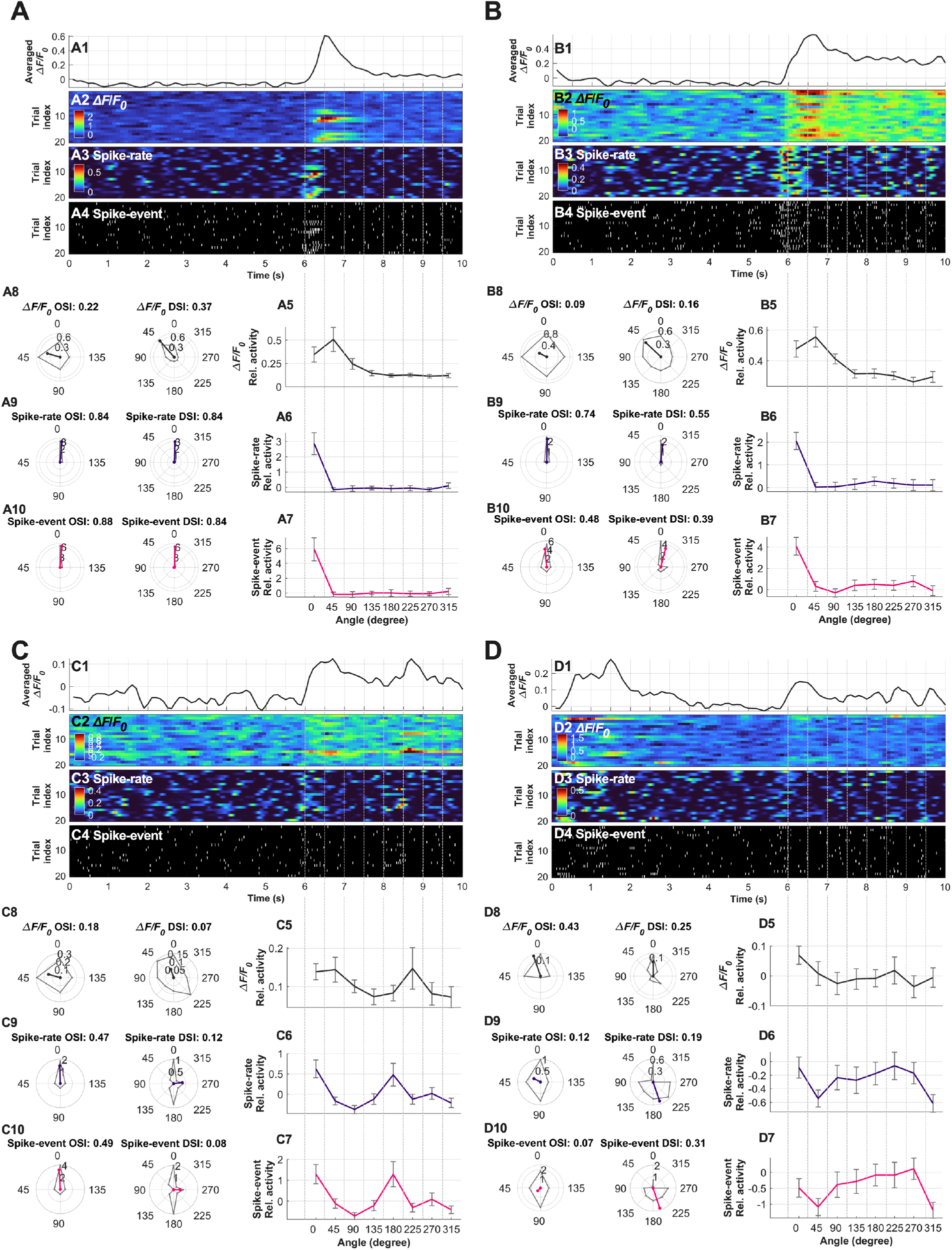
Examples of applying the ENS^2^ system to information encoding in primary visual cortex (Extension of Figure 6). Examples of recorded calcium signals and the predicted spike-rates and spike-events by ENS^2^ (**A1**-**A4, B1**-**B4, C1**-**C4, D1**-**D4**). Tuning curves and selectivity indexes (**A5**-**A10, B5**-**B10, C5**-**C10, D5**-**D10**) are computed based on three types of inputs (**A2**-**A4, B2**-**B4, C2**-**C4, D2**-**D4**), respectively. Conventions are same as **Figure 6**. Details are discussed in Results section.

**Supplementary Figure 5.**
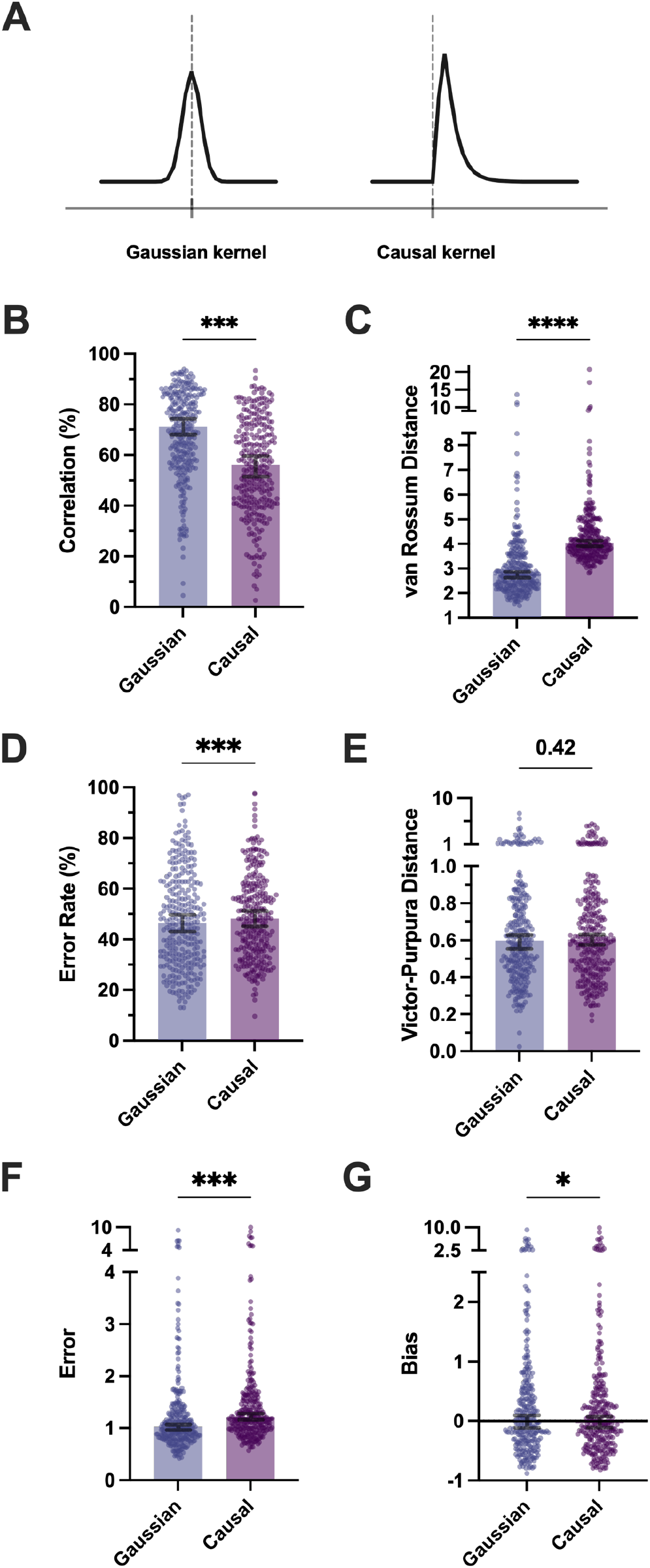
Comparison of benchmark results with ENS^2^ using different smoothing kernels. (**A**) Illustration of the resultant spike-rates when convolving one spike-event with Gaussian kernel and Causal kernel. (**B**-**G**) Comparison of performance measured by six metrics. Colored dots represent each neuron from all datasets. Statistical results are obtained by two-sided Wilcoxon signed-rank test (see Methods, ^**^p<0.01, ^***^p<0.001, ^****^p<0.0001).

**Supplementary Figure 6.**
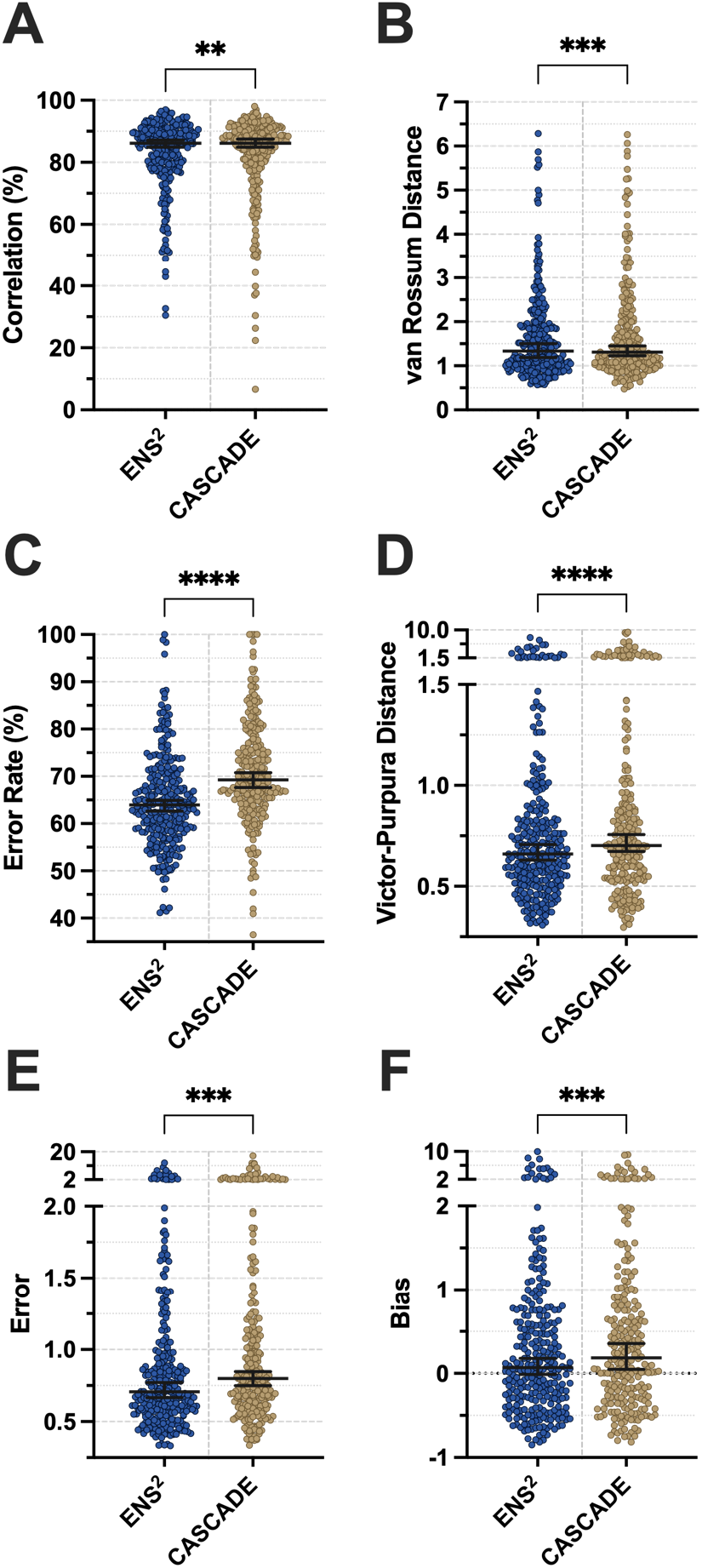
Benchmark results under 7.5Hz sampling rate with 200ms smoothing window. Our proposed ENS^2^ system shows advantages in all measuring metrics under the evaluation scheme proposed by CASCADE. Conventions are same as **Figure 3**. Statistical results are obtained by two-sided Wilcoxon signed-rank test (see Methods, ^**^p<0.01, ^***^p<0.001, ^****^p<0.0001).

**Supplementary Figure 7.**
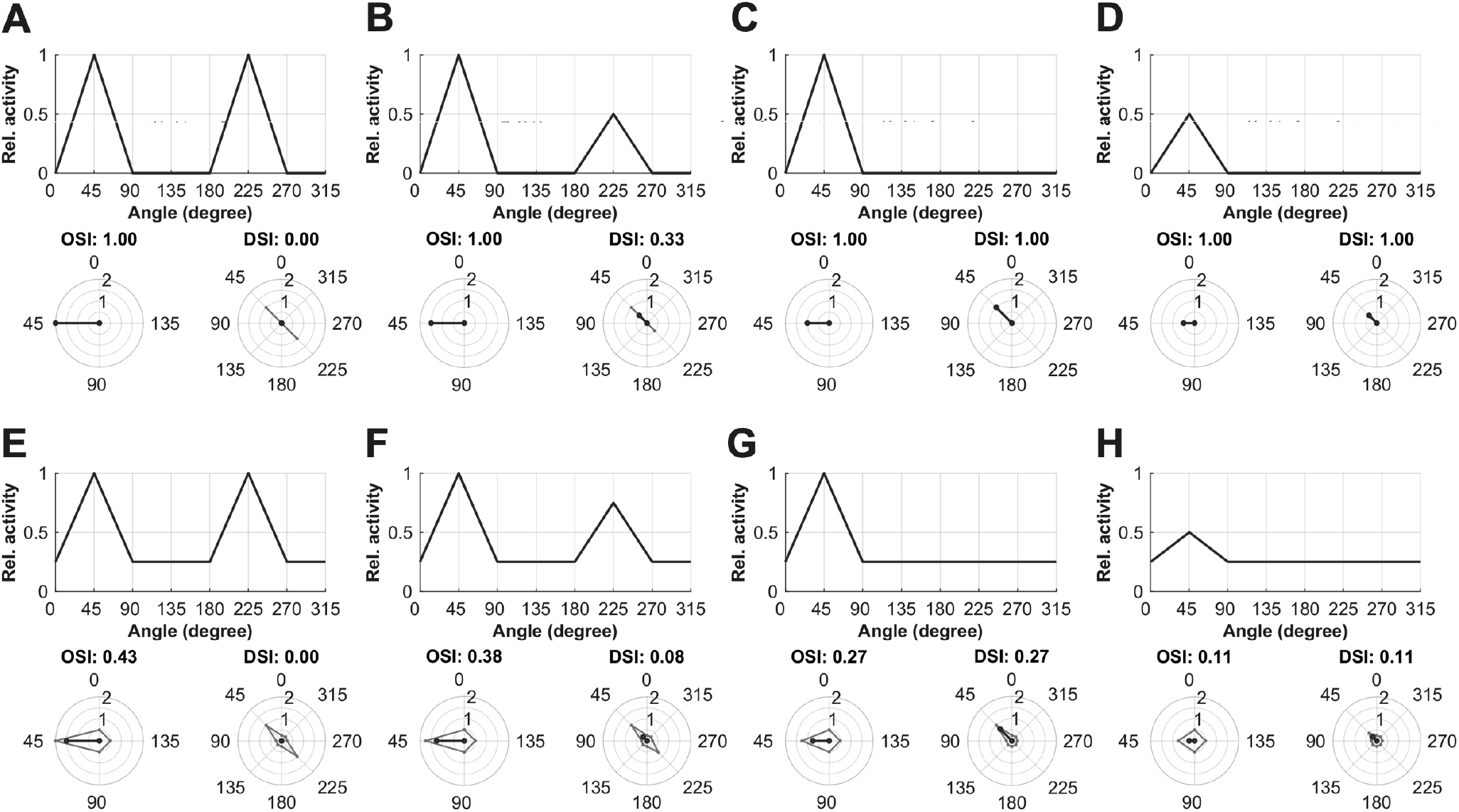
Examples of tuning curves and the corresponding OSI/DSI. Eight representative examples of tuning curves are presented as references for **Figure 6** and **Supplementary Figure 4**.

## References

Akerboom, J., Chen, T.-W., Wardill, T.J., Tian, L., Marvin, J.S., Mutlu, S., Calderón, N.C., Esposti, F., Borghuis, B.G., Sun, X.R., et al. (2012). Optimization of a GCaMP Calcium Indicator for Neural Activity Imaging. The Journal of Neuroscience 32, 13819. 10.1523/JNEUROSCI.2601-12.2012.

Alcaraz, J.M.L., and Strodthoff, N. (2022). Diffusion-based Time Series Imputation and Forecasting with Structured State Space Models. arXiv preprint 2208.09399.

Berens, P., Freeman, J., Deneux, T., Chenkov, N., McColgan, T., Speiser, A., Macke, J.H., Turaga, S.C., Mineault, P., Rupprecht, P., et al. (2018). Community-based benchmarking improves spike rate inference from two-photon calcium imaging data. PLoS Comput. Biol. 14, e1006157. 10.1371/journal.pcbi.1006157.

Bethge, P., Carta, S., Lorenzo, D.A., Egolf, L., Goniotaki, D., Madisen, L., Voigt, F.F., Chen, J.L., Schneider, B., Ohkura, M., et al. (2017). An R-CaMP1.07 reporter mouse for cell-type-specific expression of a sensitive red fluorescent calcium indicator. PLoS One 12, e0179460. 10.1371/journal.pone.0179460.

Buzsáki, G. (2004). Large-scale recording of neuronal ensembles. Nat. Neurosci. 7, 446–451. 10.1038/nn1233.

Charles, A.S., Song, A., Gauthier, J.L., Pillow, J.W., and Tank, D.W. (2019). Neural Anatomy and Optical Microscopy (NAOMi) Simulation for evaluating calcium imaging methods. bioRxiv, 726174. 10.1101/726174.

Chen, T.-W., Wardill, T.J., Sun, Y., Pulver, S.R., Renninger, S.L., Baohan, A., Schreiter, E.R., Kerr, R.A., Orger, M.B., Jayaraman, V., et al. (2013). Ultrasensitive fluorescent proteins for imaging neuronal activity. Nature 499, 295–300. 10.1038/nature12354.

Dana, H., Mohar, B., Sun, Y., Narayan, S., Gordus, A., Hasseman, J.P., Tsegaye, G., Holt, G.T., Hu, A., Walpita, D., et al. (2016). Sensitive red protein calcium indicators for imaging neural activity. eLife 5, e12727. 10.7554/eLife.12727.

de Vries, S.E.J., Lecoq, J.A., Buice, M.A., Groblewski, P.A., Ocker, G.K., Oliver, M., Feng, D., Cain, N., Ledochowitsch, P., Millman, D., et al. (2020). A large-scale standardized physiological survey reveals functional organization of the mouse visual cortex. Nat. Neurosci. 23, 138–151. 10.1038/s41593-019-0550-9.

Deneux, T., Kaszas, A., Szalay, G., Katona, G., Lakner, T., Grinvald, A., Rózsa, B., and Vanzetta, I. (2016). Accurate spike estimation from noisy calcium signals for ultrafast three-dimensional imaging of large neuronal populations in vivo. Nat. Commun. 7, 12190. 10.1038/ncomms12190.

Denk, W., Strickler, J.H., and Webb, W.W. (1990). Two-photon laser scanning fluorescence microscopy. Science 248, 73. 10.1126/science.2321027.

El-Boustani, S., Ip Jacque, P.K., Breton-Provencher, V., Knott Graham, W., Okuno, H., Bito, H., and Sur, M. (2018). Locally coordinated synaptic plasticity of visual cortex neurons in vivo. Science 360, 1349–1354. 10.1126/science.aao0862.

Éltes, T., Szoboszlay, M., Kerti-Szigeti, K., and Nusser, Z. (2019). Improved spike inference accuracy by estimating the peak amplitude of unitary [Ca2+] transients in weakly GCaMP6f-expressing hippocampal pyramidal cells. The Journal of Physiology 597, 2925–2947. 10.1113/jp277681.

Friedrich, J., and Paninski, L. (2016). Fast active set methods for online spike inference from calcium imaging. Advances In Neural Information Processing Systems 29.

Friedrich, J., Zhou, P., and Paninski, L. (2017). Fast online deconvolution of calcium imaging data. PLoS Comput. Biol. 13, e1005423. 10.1371/journal.pcbi.1005423.

Giovannucci, A., Badura, A., Deverett, B., Najafi, F., Pereira, T.D., Gao, Z., Ozden, I., Kloth, A.D., Pnevmatikakis, E., Paninski, L., et al. (2017). Cerebellar granule cells acquire a widespread predictive feedback signal during motor learning. Nat. Neurosci. 20, 727. 10.1038/nn.4531.

Giovannucci, A., Friedrich, J., Gunn, P., Kalfon, J., Brown, B.L., Koay, S.A., Taxidis, J., Najafi, F., Gauthier, J.L., Zhou, P., et al. (2019). CaImAn an open source tool for scalable calcium imaging data analysis. eLife 8, e38173. 10.7554/eLife.38173.

Goodfellow, I.J., Pouget-Abadie, J., Mirza, M., Xu, B., Warde-Farley, D., Ozair, S., Courville, A., and Bengio, Y. (2014). Generative adversarial nets. Proceedings of the 27th International Conference on Neural Information Processing Systems - Volume 2. MIT Press.

Greenberg, D.S., Houweling, A.R., and Kerr, J.N.D. (2008). Population imaging of ongoing neuronal activity in the visual cortex of awake rats. Nat. Neurosci. 11, 749–751. 10.1038/nn.2140.

Grewe, B.F., Langer, D., Kasper, H., Kampa, B.M., and Helmchen, F. (2010). High-speed in vivo calcium imaging reveals neuronal network activity with near-millisecond precision. Nat. Methods 7, 399–405. 10.1038/nmeth.1453.

Grienberger, C., and Konnerth, A. (2012). Imaging Calcium in Neurons. Neuron 73, 862–885. 10.1016/j.neuron.2012.02.011.

Hamill, O.P., Marty, A., Neher, E., Sakmann, B., and Sigworth, F.J. (1981). Improved patch-clamp techniques for high-resolution current recording from cells and cell-free membrane patches. Pflügers Archiv 391, 85–100. 10.1007/BF00656997.

Ho, J., Jain, A., and Abbeel, P. (2020). Denoising diffusion probabilistic models. Advances in Neural Information Processing Systems 33, 6840–6851.

Hoang, H., Sato, M.-a., Shinomoto, S., Tsutsumi, S., Hashizume, M., Ishikawa, T., Kano, M., Ikegaya, Y., Kitamura, K., Kawato, M., and Toyama, K. (2020). Improved hyperacuity estimation of spike timing from calcium imaging. Sci. Rep. 10, 17844. 10.1038/s41598-020-74672-y.

Huang, L., Knoblich, U., Ledochowitsch, P., Lecoq, J., Reid, R.C., de Vries, S.E.J., Buice, M.A., Murphy, G.J., Waters, J., Koch, C., et al. (2020). Relationship between simultaneously recorded spiking activity and fluorescence signal in GCaMP6 transgenic mice. bioRxiv, 788802. 10.1101/788802.

Jewell, S.W., Hocking, T.D., Fearnhead, P., and Witten, D.M. (2020). Fast nonconvex deconvolution of calcium imaging data. Biostatistics 21, 709–726. 10.1093/biostatistics/kxy083.

Kerr, J.N.D., and Denk, W. (2008). Imaging in vivo: watching the brain in action. Nat. Rev. Neurosci. 9, 195–205. 10.1038/nrn2338.

Kerr, J.N.D., Greenberg, D., and Helmchen, F. (2005). Imaging input and output of neocortical networks in vivo. Proceedings of the National Academy of Sciences of the United States of America 102, 14063. 10.1073/pnas.0506029102.

Khan, A.G., Poort, J., Chadwick, A., Blot, A., Sahani, M., Mrsic-Flogel, T.D., and Hofer, S.B. (2018). Distinct learning-induced changes in stimulus selectivity and interactions of GABAergic interneuron classes in visual cortex. Nat. Neurosci. 21, 851–859. 10.1038/s41593-018-0143-z.

Kingma, D.P., and Ba, J. (2015). Adam: a method for stochastic optimization. 3rd International Conference on Learning Representations (ICLR).

Knogler, L.D., Markov, D.A., Dragomir, E.I., Štih, V., and Portugues, R. (2017). Sensorimotor Representations in Cerebellar Granule Cells in Larval Zebrafish Are Dense, Spatially Organized, and Non-temporally Patterned. Current Biology 27, 1288–1302. 10.1016/j.cub.2017.03.029.

Kwan, Alex C., and Dan, Y. (2012). Dissection of Cortical Microcircuits by Single-Neuron Stimulation In Vivo. Current Biology 22, 1459–1467.

Lecun, Y., Bottou, L., Bengio, Y., and Haffner, P. (1998). Gradient-based learning applied to document recognition. Proc. IEEE 86, 2278–2324. 10.1109/5.726791.

Li, X., Zhang, G., Wu, J., Zhang, Y., Zhao, Z., Lin, X., Qiao, H., Xie, H., Wang, H., Fang, L., and Dai, Q. (2020). Reinforcing neuron extraction and spike inference in calcium imaging using deep self-supervised learning. bioRxiv, 2020.2011.2016.383984. 10.1101/2020.11.16.383984.

Mazurek, M., Kager, M., and Van Hooser, S.D. (2014). Robust quantification of orientation selectivity and direction selectivity. Front. Neural Circuits 8, 92–92. 10.3389/fncir.2014.00092.

Neher, E., and Sakmann, B. (1976). Single-channel currents recorded from membrane of denervated frog muscle fibres. Nature 260, 799–802. 10.1038/260799a0.

Onativia, J., Schultz, S.R., and Dragotti, P.L. (2013). A finite rate of innovation algorithm for fast and accurate spike detection from two-photon calcium imaging. J. Neural Eng. 10, 046017.

Pachitariu, M., Stringer, C., Dipoppa, M., Schröder, S., Rossi, L.F., Dalgleish, H., Carandini, M., and Harris, K.D. (2017). Suite2p: beyond 10,000 neurons with standard two-photon microscopy. bioRxiv, 061507. 10.1101/061507.

Pachitariu, M., Stringer, C., and Harris, K.D. (2018). Robustness of Spike Deconvolution for Neuronal Calcium Imaging. The Journal of Neuroscience 38, 7976. 10.1523/JNEUROSCI.3339-17.2018.

Pnevmatikakis, E.A. (2019). Analysis pipelines for calcium imaging data. Curr. Opin. Neurobiol. 55, 15–21.

Pnevmatikakis, E.A., Merel, J., Pakman, A., and Paninski, L. (2013). Bayesian spike inference from calcium imaging data. 3-6 Nov. 2013. pp. 349–353.

Pnevmatikakis, Eftychios A., Soudry, D., Gao, Y., Machado, T.A., Merel, J., Pfau, D., Reardon, T., Mu, Y., Lacefield, C., Yang, W., et al. (2016). Simultaneous Denoising, Deconvolution, and Demixing of Calcium Imaging Data. Neuron 89, 285–299. 10.1016/j.neuron.2015.11.037.

Rahmati, V., Kirmse, K., Marković, D., Holthoff, K., and Kiebel, S.J. (2016). Inferring Neuronal Dynamics from Calcium Imaging Data Using Biophysical Models and Bayesian Inference. PLoS Comput. Biol. 12, e1004736. 10.1371/journal.pcbi.1004736.

Rikhye, R.V., and Sur, M. (2015). Spatial Correlations in Natural Scenes Modulate Response Reliability in Mouse Visual Cortex. The Journal of Neuroscience 35, 14661. 10.1523/JNEUROSCI.1660-15.2015.

Ronneberger, O., Fischer, P., and Brox, T. (2015). U-Net: convolutional networks for biomedical image segmentation. held in Cham, N. Navab, J. Hornegger, W.M. Wells, and A.F. Frangi, eds. (Springer International Publishing), pp. 234–241.

Rupprecht, P., Carta, S., Hoffmann, A., Echizen, M., Blot, A., Kwan, A.C., Dan, Y., Hofer, S.B., Kitamura, K., Helmchen, F., and Friedrich, R.W. (2021). A database and deep learning toolbox for noise-optimized, generalized spike inference from calcium imaging. Nat. Neurosci. 10.1038/s41593-021-00895-5.

Schoenfeld, G., Carta, S., Rupprecht, P., Ayaz, A., and Helmchen, F. (2021). In vivo calcium imaging of CA3 pyramidal neuron populations in adult mouse hippocampus. bioRxiv, 2021.2001.2021.427642. 10.1101/2021.01.21.427642.

Sebastian, J., Kumar, M.G., Viraraghavan, V.S., Sur, M., and Murthy, H.A. (2019). Spike Estimation From Fluorescence Signals Using High-Resolution Property of Group Delay. IEEE Trans. Signal Process. 67, 2923–2936. 10.1109/TSP.2019.2908913.

Sebastian, J., Sur, M., Murthy, H.A., and Magimai-Doss, M. (2021). Signal-to-signal neural networks for improved spike estimation from calcium imaging data. PLoS Comput. Biol. 17, e1007921. 10.1371/journal.pcbi.1007921.

Sofroniew, N.J., Flickinger, D., King, J., and Svoboda, K. (2016). A large field of view two-photon mesoscope with subcellular resolution for in vivo imaging. eLife 5, e14472. 10.7554/eLife.14472.

Spira, M.E., and Hai, A. (2013). Multi-electrode array technologies for neuroscience and cardiology. Nat. Nanotechnol. 8, 83–94. 10.1038/nnano.2012.265.

Srivastava, N., Hinton, G., Krizhevsky, A., Sutskever, I., and Salakhutdinov, R. (2014). Dropout: a simple way to prevent neural networks from overfitting. The journal of machine learning research 15, 1929–1958.

Stosiek, C., Garaschuk, O., Holthoff, K., and Konnerth, A. (2003). In vivo two-photon calcium imaging of neuronal networks. Proceedings of the National Academy of Sciences 100, 7319. 10.1073/pnas.1232232100.

Stringer, C., and Pachitariu, M. (2019). Computational processing of neural recordings from calcium imaging data. Curr. Opin. Neurobiol. 55, 22–31. 10.1016/j.conb.2018.11.005.

Tada, M., Takeuchi, A., Hashizume, M., Kitamura, K., and Kano, M. (2014). A highly sensitive fluorescent indicator dye for calcium imaging of neural activity in vitro and in vivo. European Journal of Neuroscience 39, 1720–1728. 10.1111/ejn.12476.

Tashiro, Y., Song, J., Song, Y., and Ermon, S. (2021). CSDI: Conditional score-based diffusion models for probabilistic time series imputation. Advances in Neural Information Processing Systems 34, 24804–24816.

Theis, L., Berens, P., Froudarakis, E., Reimer, J., Román Rosón, M., Baden, T., Euler, T., Tolias Andreas S., and Bethge, M. (2016). Benchmarking Spike Rate Inference in Population Calcium Imaging. Neuron 90, 471–482. 10.1016/j.neuron.2016.04.014.

Tsutsumi, S., Yamazaki, M., Miyazaki, T., Watanabe, M., Sakimura, K., Kano, M., and Kitamura, K. (2015). Structure-function relationships between aldolase C/zebrin II expression and complex spike synchrony in the cerebellum. The Journal of neuroscience: the official journal of the Society for Neuroscience 35, 843–852. 10.1523/JNEUROSCI.2170-14.2015.

Ulyanov, D., Vedaldi, A., and Lempitsky, V. (2016). Instance normalization: The missing ingredient for fast stylization. arXiv preprint 1607.08022.

van Rossum, M.C. (2001). A novel spike distance. Neural Comput 13, 751–763. 10.1162/089976601300014321.

Vaswani, A., Shazeer, N., Parmar, N., Uszkoreit, J., Jones, L., Gomez, A.N., Kaiser, L., and Polosukhin, I. (2017). Attention Is All You Need. arXiv e-prints, 1706.03762.

Victor, J.D., and Purpura, K.P. (1996). Nature and precision of temporal coding in visual cortex: a metric-space analysis. Journal of Neurophysiology 76, 1310–1326. 10.1152/jn.1996.76.2.1310.

Vogelstein, J.T., Packer, A.M., Machado, T.A., Sippy, T., Babadi, B., Yuste, R., and Paninski, L. (2010). Fast Nonnegative Deconvolution for Spike Train Inference From Population Calcium Imaging. Journal of Neurophysiology 104, 3691–3704. 10.1152/jn.01073.2009.

Wagner, M.J., Kim, T.H., Savall, J., Schnitzer, M.J., and Luo, L. (2017). Cerebellar granule cells encode the expectation of reward. Nature 544, 96. 10.1038/nature21726.

Wilson, N.R., Runyan, C.A., Wang, F.L., and Sur, M. (2012). Division and subtraction by distinct cortical inhibitory networks in vivo. Nature 488, 343–348. 10.1038/nature11347.

Yaksi, E., and Friedrich, R.W. (2006). Reconstruction of firing rate changes across neuronal populations by temporally deconvolved Ca2+ imaging. Nat. Methods 3, 377–383. 10.1038/nmeth874.

Zhai, X., Jelfs, B., Chan, R.H.M., and Tin, C. (2017). Self-recalibrating surface EMG pattern recognition for neuroprosthesis control based on convolutional neural network. Front. Neurosci. 11. 10.3389/fnins.2017.00379.

Zhai, X., and Tin, C. (2018). Automated ECG classification using dual heartbeat coupling based on convolutional neural network. IEEE Access 6, 27465–27472. 10.1109/access.2018.2833841.

Zhai, X., Zhou, Z., and Tin, C. (2020). Semi-supervised learning for ECG classification without patient-specific labeled data. Expert Syst. Appl. 158, 113411. 10.1016/j.eswa.2020.113411.

Zhou, Z., Zhai, X., and Tin, C. (2021). Fully automatic electrocardiogram classification system based on generative adversarial network with auxiliary classifier. Expert Syst. Appl. 174, 114809. 10.1016/j.eswa.2021.114809.

